# In vivo high-content screening in zebrafish for developmental nephrotoxicity of approved drugs

**DOI:** 10.1101/2020.04.21.052688

**Authors:** JH Westhoff, PJ Steenbergen, LSV Thomas, J Heigwer, T Bruckner, L Cooper, B Tönshoff, GF Hoffmann, J Gehrig

**Author notes:** Institute of Cancer and Genomic Sciences, College of Medical and Dental Sciences, University of Birmingham, Birmingham, United Kingdom (L Cooper).

## Abstract

Despite widespread drug exposure, for example during gestation or in prematurely born children, organ-specific developmental toxicity of most drugs is poorly understood. Developmental and functional abnormalities are a major cause of kidney diseases during childhood; however, the potential causal relationship to exposure with nephrotoxic drugs during nephrogenesis is widely unknown. To identify developmental nephrotoxic drugs in a large scale, we established and performed an automated high-content screen to score for phenotypic renal alterations in the *Tg(wt1b:EGFP)* zebrafish line. During early nephrogenesis, embryos were exposed to a compound library of approved drugs. After treatment, embryos were aligned within microtiter plates using 3D-printed orientation tools enabling the robust acquisition of consistent dorsal views of pronephric kidneys by automated microscopy. To qualitatively and quantitatively score and visualize phenotypes, we developed software tools for the semi-automated analysis, processing and visualization of this large image-based dataset. Using this scoring scheme, we were able to categorize compounds based on their potential developmental nephrotoxic effects. About 10% of tested drugs induced pronephric phenotypes including glomerular and tubular malformations, or overall changes in kidney morphology. Major chemical compound groups identified to cause glomerular and tubular alterations included dihydropyridine derivatives, HMG CoA reductase inhibitors, fibrates, imidazole, benzimidazole and triazole derivatives, corticosteroids, glucocorticoids, acetic acid derivatives and propionic acid derivatives. In conclusion, the presented study demonstrates the large-scale screening of kidney-specific toxicity of approved drugs in a live vertebrate embryo. The associated technology and tool-sets can be easily adapted for other organ systems providing a unique platform for *in vivo* large-scale assessment of organ-specific developmental toxicity or other biomedical applications. Ultimately, the presented data and associated visualization and browsing tools provide a resource for potentially nephrotoxic drugs and for further investigations.

## Introduction

Pharmaceutical drugs and other chemicals can negatively impact organogenesis, either during pregnancy or by postnatal exposure of very preterm infants. It has been estimated that up to 10% of congenital anomalies may be caused by environmental exposures including adverse drug effects (Brent, 2004). Mostly unaware of pregnancy, 22-50% of women take medication in the first trimester (De Vigan et al., 1999). During the entire pregnancy, 65-94% of women get at least one drug prescription (Ramoz and Patel-Shori, 2014) with the use of over-the-counter medications being assumed even higher (Werler et al., 2005). However, safety data on drug-induced organ-specific developmental toxicity of chemicals registered for commercial use is scarce and often solely based on observational human studies and case reports.

Developmental toxicity testing of promising drug candidates according to current international guidelines includes a series of screening tests (e.g. OECD Prenatal Developmental Toxicity Study, Two-Generation Reproduction Toxicity Study, Reproduction/Developmental Toxicity Screening Test) that are predominantly performed in historically established mammalian models (e.g. rat, rabbit). However, these *in vivo* tests require large numbers of animals, thus raising ethical concerns, and are highly laborious and expensive (Piersma, 2004).

Besides classic *in vitro* tests lacking physiological context, *in vivo* high-throughput screening assays using small model organisms have been proposed, which have the potential to bridge the gap between cell-based assays and mammalian animal models (Piersma, 2004). The cost-effective and readily accessible zebrafish has been established as a main vertebrate model for human disease modeling, drug discovery and safety applications as well as toxicological studies (Brady et al., 2016, MacRae and Peterson, 2015). Small size, *ex utero* development, optical transparency, and rapidity of organogenesis render zebrafish embryos and larvae ideal for *in vivo* high-content screening applications and large-scale assessment of compound effects. The increasing usage in large-scale assays is further driven by efforts for more ethical use of animals in testing, as under current legislation zebrafish embryos and larvae comply with the 3R principles (Replacement, Reduction and Refinement) (Ball et al., 2014, Kirk, 2018, Russell, 1995). Moreover, the zebrafish genome harbors orthologues of 70% of human genes including 86% of known drug targets, underscoring its potential to identify teratogenicity hazard (Gunnarsson et al., 2008, Howe et al., 2013). Importantly, the concordance between zebrafish and mammalian developmental toxicity has been evaluated in several studies and may be higher than 80% (Brannen et al., 2010, Nagel, 2002).

Nephrogenesis in humans continues until about 34–36 weeks of gestation with more than 60% of nephrons being formed in the last trimester of pregnancy (Mackenzie and Brenner, 1995, Rodriguez et al., 2004). Congenital anomalies of the kidney including renal hypoplasia and dysplasia, representing the most frequent etiologies for childhood chronic kidney disease, are attributed, besides genetic factors, to adverse fetal environmental factors that include exposure to developmental nephrotoxic drugs. In addition, prematurity and low birth-weight *per se* lead to an impaired nephrogenesis, to low nephron endowment and an increased risk of neonatal acute kidney injury and chronic kidney disease (Perico et al., 2018). Strikingly, long-term studies on the nephrotoxic action of drugs administered during active nephrogenesis are mostly lacking. Still, several acute toxic and teratogenic effects on human kidney development have been described, amongst others, for angiotensin-converting enzyme (ACE) inhibitors (Mastrobattista, 1997), angiotensin type 1 (AT1) receptor antagonists (Hinsberger et al., 2001, Payen et al., 2006, Boubred et al., 2006) and nonsteroidal anti-inflammatory drugs (NSAID) (Boubred et al., 2006).

Despite being a member of the teleost family, there are high similarities at the developmental and cell- and organ-specific level between zebrafish and humans (19). The pronephros of the zebrafish larva consists of two nephrons with a fused glomerulus ventrally to the dorsal aorta. While tremendous differences in nephron number between humans and zebrafish larvae exist, their homology results in a highly similar composition of single nephrons at the cellular and molecular level (Drummond, 2005, Drummond and Davidson, 2010). In addition, kidney development and function largely depend on the same orthologous genes for all vertebrate kidneys. Therefore, studying the impact of compounds on zebrafish pronephros formation and function can aid in the understanding of drug-induced harmful effects on renal development in humans (Drummond, 2005).

To score developmental phenotypes of the zebrafish developing kidney in a large scale, we have previously reported the development of an automated imaging pipeline for the analysis of pronephroi in live zebrafish embryos and larvae (Westhoff et al., 2013). In a pilot screen, we could show that human developmental nephrotoxic drugs also induced morphological alterations in developing larval zebrafish pronephroi. Here, we improved this screening assay and technology, and performed a large *in vivo* high-content screening experiment with the aim to identify approved drugs with adverse effects on pronephros development in zebrafish. We present the results of this large-scale whole organism screen obtained by scoring multiple qualitative and quantitative morphological phenotypic parameters in live transgenic zebrafish embryos. This revealed several compound classes, with partially similar chemical properties or mode of action that negatively impact nephrogenesis in the zebrafish model system.

## Results

### Screening assay for developmental nephrotoxicity

To establish a screening protocol and microscopy workflow compatible with the required handling and phenotypic scoring of thousands of compound-treated transgenic zebrafish embryos, we expanded and refined our previously published high-content screening pipeline for automated imaging of standardized dorsal views of the developing pronephros in zebrafish embryos (**Figure 1**) (Westhoff et al., 2013). In brief, embryos of the *Tg(wt1b:EGFP*) transgenic line were enzymatically dechorionated at 24 hours post fertilization (hpf), and then exposed to 1,280 drugs of the Prestwick Chemical Library® at a concentration of 25 µM for a 24 hour period (**Figure 1A**). This concentration was chosen based on pilot experiments with known nephrotoxic drugs, which aimed to identify concentration ranges causing clear renal pathological phenotypes while keeping general toxicity low (data not shown). A total of 20 embryos were exposed to each compound. At 48 hpf, brightfield overview images were taken using a stereo microscope and gross morphological alterations were evaluated (**Supplementary Tables 1, 2**). Subsequently, automated acquisition of stable and consistent dorsal views of pronephric kidneys (n = 7-12 larvae, depending on lethality rates) were acquired (**Figure 1B**). This was enabled by usage of 3D-printed orientation tools to create cavities in agarose-filled microplates that enable consistent positioning and dorsal orientation of specimen (Wittbrodt et al., 2014). To ensure capturing of entire organs and compensate for minor variations in z-positioning, each larva was imaged using z-stacks with 10 slices (Δz=15 µM) in the brightfield and GFP channel. In total, >15,000 compound treated embryos represented in >300,000 images were acquired. The orientation tool permits organ and tissue specific screening on standard screening microscopes using a fixed field of view for all wells using lower magnification objectives, resulting in image data at sufficient resolution to score overall morphological phenotypes of the pronephros (Westhoff et al., 2013, Wittbrodt et al., 2014). To adequately visualize more delicate features, such as boundaries of glomerular cysts, software modules for automatic detection and centering the region of interests are recommended (Pandey et al., 2019). Automated data acquisition of z-stacks was followed by file handling including sorting and generation of multilayer tiff files. To enable rapid visual assessment of screening results, image data processing was carried out to generate maximum z-projection images followed by automated region of interest detection, cropping of kidney regions and generation of thumbnail overview images (**Figure 1B**).

**Figure 1.**
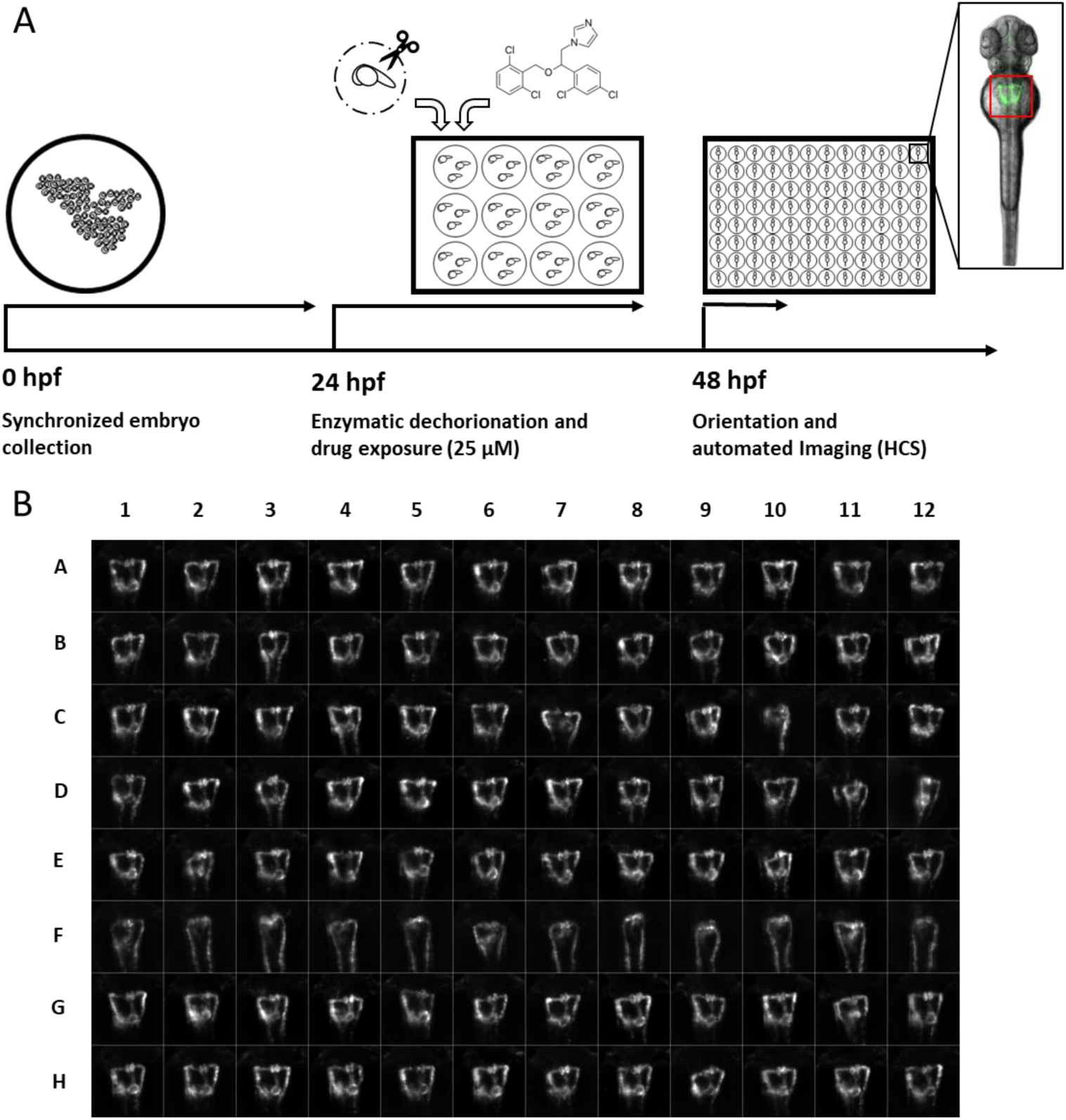
A screening workflow to score potential nephrotoxicity of approved drugs. (**A**) Schematic illustration of the drug toxicity screening pipeline. Synchronized embryos were obtained by collecting eggs from batch and pairwise crossings within a maximum interval of 30 minutes. At 24 hours post fertilization (hpf) green fluorescent protein (GFP) positive embryos were enzymatically dechorionated using Pronase and 20 embryos were transferred in each well of a 12-well plate. Different compounds were added to each well to a final concentration of 25 µM. At 48 hpf, and after 24 hour drug exposure, embryos were washed, transferred into agarose filled microtiter plates, oriented within cavities generated with 3D-printed orientation tools and automatically imaged. The location of embryonic kidneys within image data was automatically determined using center of mass detection after thresholding of maximum z-projections. Center of mass coordinates were used to automatically crop maximum z-projections to thumbnails images with 257 × 257 pixels dimension (see also (B)). Thumbnail images were used for all subsequent analysis steps. (**B**) Representative example of montage of pronephros thumbnail images, which was generated for each experimental 96-well plate. Each row was loaded with differently treated embryos. Row A-G show compound treated embryos, row F shows embryos with severely altered pronephros morphology, and row H shows embryos treated only with DMSO serving as plate internal negative controls.

### Renal and non-renal scoring parameters

Kidney morphology as imaged by automated fluorescence microscopy was scored quantitatively and qualitatively. Quantitative analysis was performed using a Fiji macro that automatically calculated certain tissue dimensions and parameters after manual definition of 16 reference points (Schindelin et al., 2012) (**Figure 2A** and **Supplementary Software 1**). This included measurement of the following tubular structures: i) maximum distance between the tubules, ii) left and right angle between neck segment and proximal convoluted tubule, and iii) left and right proximal tubular diameter. Glomerular parameters analyzed included i) glomerular height and ii) glomerular width as markers for glomerular malformation, and iii) the distance between the glomeruli (**Figure 2B**). The second approach to evaluate kidney morphology included the development of a software annotation tool that permitted blinded manual assessment of renal phenotypes by assigning categories or image tags to random images of a predefined selection by an expert annotator (**Figure 2C**). Qualitative parameters included reduced pronephric angle (major, minor, moderate) between neck segment and proximal convoluted tubule, malformed glomeruli and glomerular separation (major, minor, moderate). The *Tg(wt1b::GFP)* transgenic line has further been shown to visualize the exocrine pancreas, which might be attributed to enhancer elements of the 5’ neighboring gene *ga17*, since *wt1b* expression cannot be detected in these tissues (Perner et al., 2007). This additional transgene activity was used to also score absence of a pancreatic signal and hence putative pancreatic damage from fluorescence z-stack images as an indicator of extrarenal toxicity. Other potentially observed phenotypic features, such as appearance of glomerular cyst, were also noted down (**Supplementary Table 1**).

**Figure 2.**
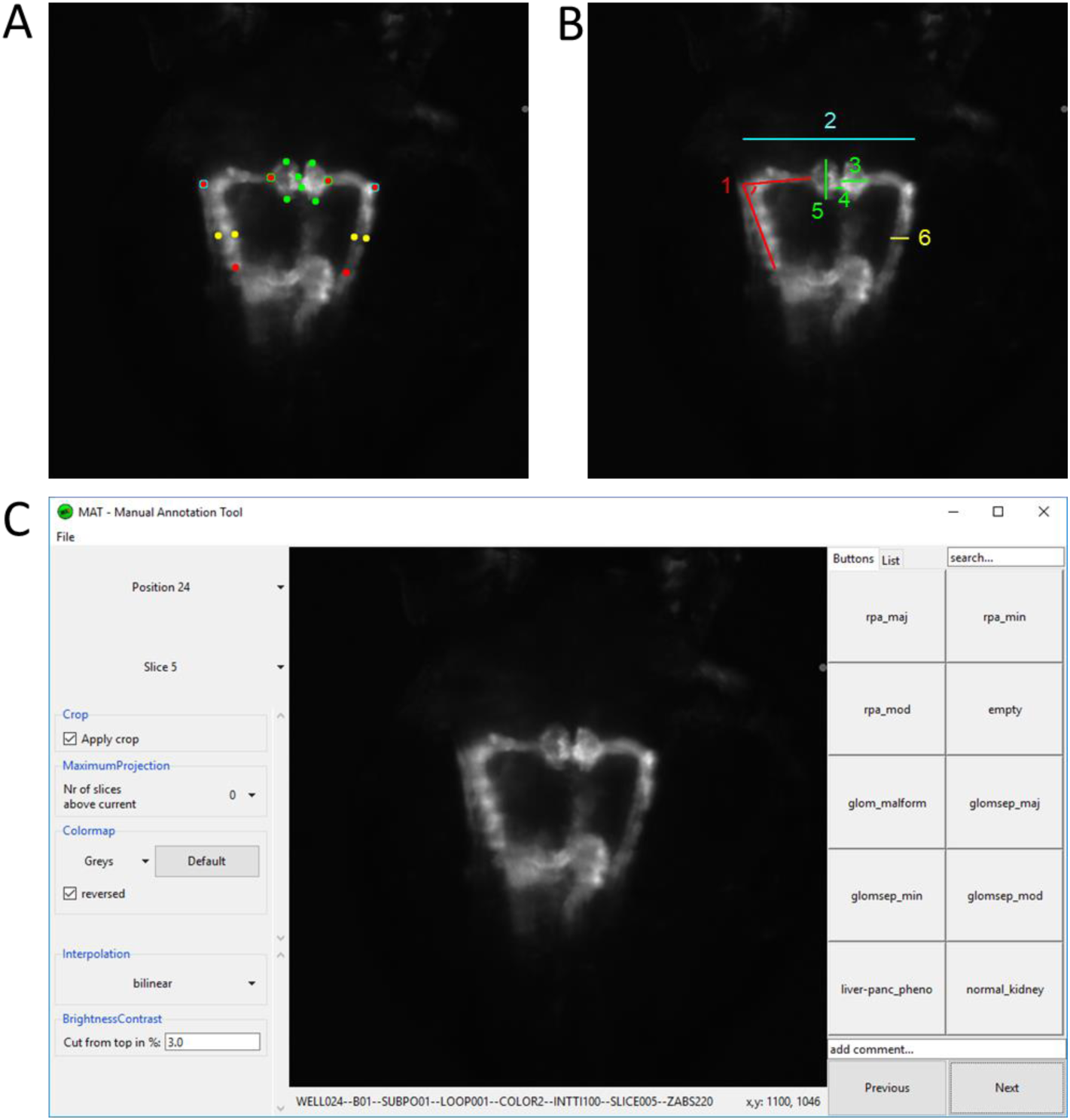
Analysis and annotation of pronephric phenotypes. (**A, B**) Quantitative measuring of morphological phenotypes of pronephroi. (A) Positions of 16 reference points are indicated that were manually assigned to each pronephros thumbnail image. (**B**) Illustration of calculated morphological parameters based on reference points in A. Colors in A and B indicate which reference points in A were used for the calculation of morphological features in B: 1) pronephric angle (only left side is shown - angleL), 2) tubular distance (tubDist), 3) glomerular height (only right side is shown - glomHeightR), 4) glomerular separation (glomSep), 5) glomerular width (only left side is shown - glomWidthL) and 6) tubular diameter (only right side is shown - tubDiamR). (**C**) Screenshot of software tool for manual annotation of large image datasets (https://doi.org/10.5281/zenodo.3367365). The manual annotation tool allows to browse, filter, auto-center, visualize and annotate complex multidimensional image datasets in an intuitive and blinded manner without any additional data pre-processing steps. In this study, the tool was used to assign up to 10 phenotypic categories to each acquired embryonic pronephros. Abbreviations: reduced pronephron angle major (rpa_maj), reduced pronephron angle minor (rpa_min), reduced pronephros angle moderate (rpa_mod), glomerular malformation (glom_malform), glomerular separation major (glomsep_maj), glomerular separation minor (glomsep_min), glomerular separation moderate (glomsep_mod), impaired liver pancreas area (liver-panc_pheno) and normal kidney (normal_kidney).

Prior to automated screening, overall general phenotypes were also assessed including the number/percentage of dead larvae, number/percentage of larvae with a curved back or tail, number/percentage of larvae showing either mild or severe edema, presence of yolk sac necrosis, malformed somites and heartbeat alterations (faster, slower, absent) (**Supplementary Figure 1** and **Supplementary Tables 1,2**).

### Renal phenotyping

The results of the quantitative measurement based on reference points were outlier filtered and normalized by pronephric measurements of associated negative controls values. This resulted in fold change ratios representing the phenotypic change for each quantitative parameter of every assayed compound. To facilitate comparative analysis, hit detection and visualization of data, these phenotypic changes were also z-score normalized. Qualitative parameters assigned using the Manual Annotation Tool (MAT) software were converted into ratios of embryos that were annotated with specific categories (**Figures 3,4** and **Supplementary Tables 1,2**). To generate a comprehensive overview visualization of the dataset, the 1,280 compounds list was sorted according the D-level of the official Anatomical Therapeutic Chemical (ATC) Classification System code (https://www.who.int/medicines/regulation/medicines-safety/toolkit_methodology/en/) and visualized using heat maps and parallel coordinates plots (**Figures 3,4**). Assignment of the Prestwick Chemical Library® compounds to this ATC-Classification generated 374 groups according to the organ or organ system they affect and their chemical, pharmacological and therapeutic properties. Of note, 207 out of 1,280 compounds could not be assigned to a specific group by use of the ATC Classification System at date of analysis, and hence are listed as N/A class (**Supplementary Table 2**). The ATC-code sorted heat map visualization allows identifying groups of compounds that altered pronephros morphology (**Figure 3A**). Interestingly, it revealed substance group-specific clusters of pathological renal phenotypes, and certain substances with chemical structure similarities (e.g. aromatic heterocyclic compounds including triazoles, imidazoles and benzimidazoles) produced comparable renal phenotypes even if drugs belonged to different ATC groups (**Figure 3B**).

**Figure 3.**
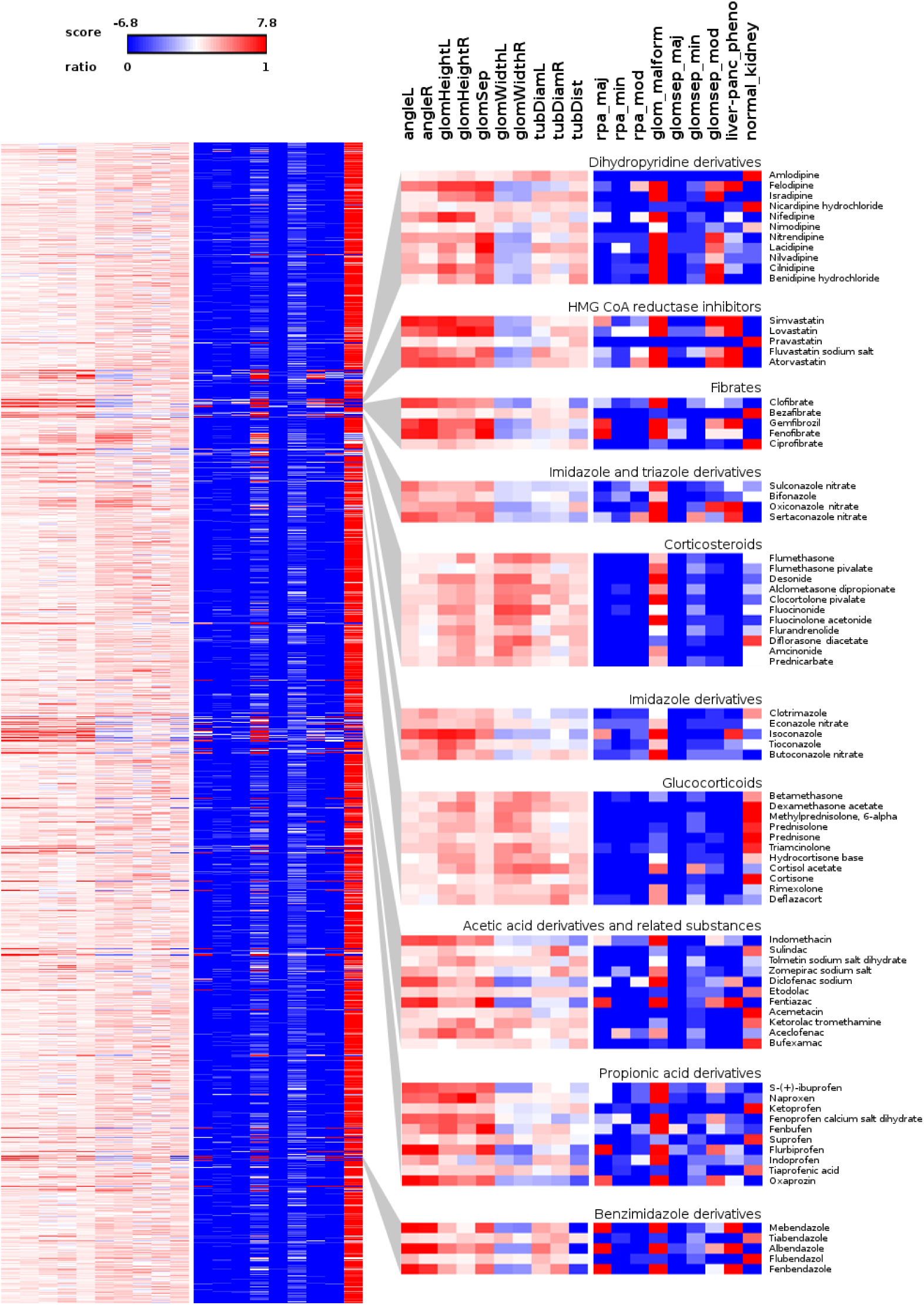
Overview of scored quantitative morphometric parameters and qualitative manual annotations. (**A**) Heat map visualizations illustrating the pronephric phenotypic alterations upon compound treatment for each assayed compound. Heat maps are sorted according to the D-level of the Anatomical Therapeutic Chemical (ATC) Classification System (for fully labeled heat maps please refer to Supplementary Figures 3 and 4). The left panel shows quantitative parameters shown as color-coded z-score. Shown are z-score changes of pronephric angle left (angleL), pronephric angle right (angleR), glomerular height left (glomHeightL), glomerular height right (glomHeightR), glomerular separation (glomSep), glomerular width left (glomWidthL), glomerular width right (glomWidthR), tubular diameter left (tubDiamL), tubular diameter right (tubDiamR) and tubular distance (tubDist). See also labelling of heat map columns in panel B. The right panel shows qualitative annotations as a ratio of embryos assigned with a certain category. Legend indicates colors assigned to values. Shown are ratios for reduced pronephron angle major (rpa_maj), reduced pronephron angle minor (rpa_min), reduced pronephros angle moderate (rpa_mod), glomerular malformation (glom_malform), glomerular separation major (glomsep_maj), glomerular separation minor (glomsep_min), glomerular separation moderate (glomsep_mod), impaired liver pancreas area (liver-panc_pheno) and normal kidney (normal_kidney). See also labelling of heat map columns in panel B. (**B**) Magnified view on illustrative examples of compound classes enriched with substances causing pronephric phenotypes.

**Figure 4.**
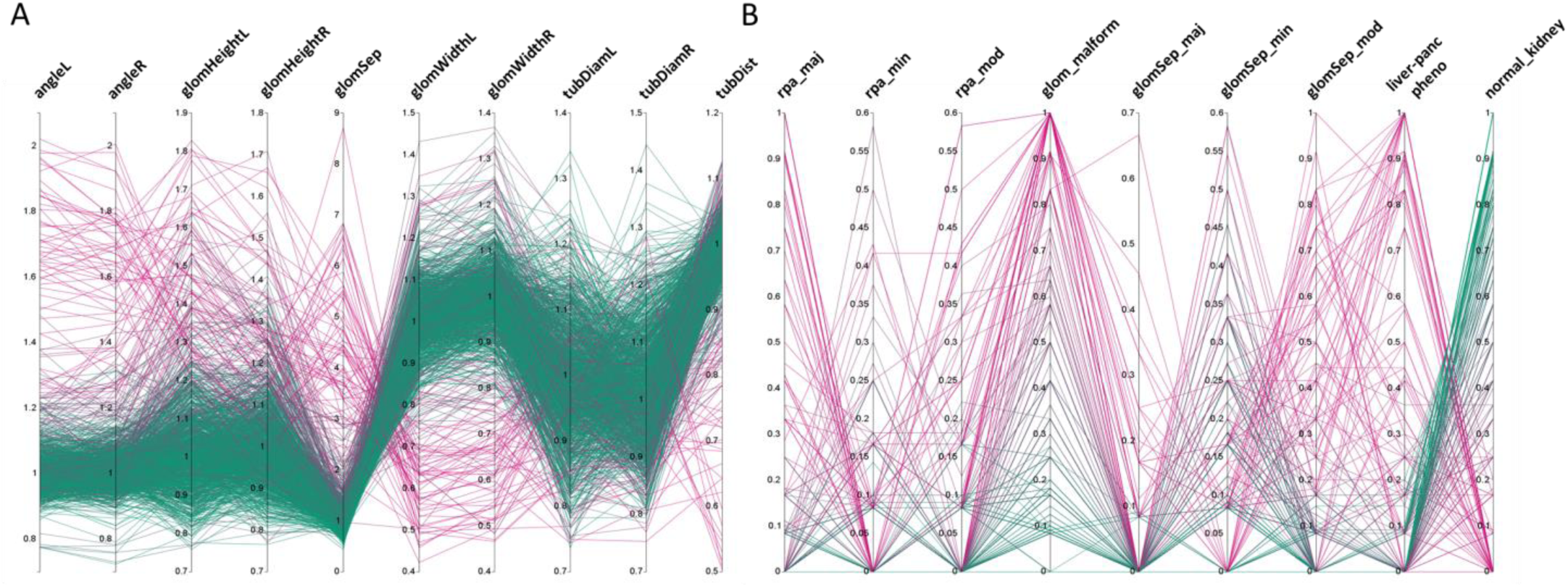
Distribution of phenotypic features upon compound treatment. Parallel coordinates plot visualizations showing the distribution of phenotypic features of pronephroi upon compound treatment. Each line plot represents a single compound treatment. In total, 1237 compound treatments are shown. The color of lines indicates the ratio of embryos within one treatment group scored as ‘normal_kidney’ using manual annotation (from 0% (magenta, all abnormal) to 100% (green, all normal)). Y-axes show the (**A**) mean fold change for different quantitative morphometric parameters, or (**B**) ratio of embryos assigned with a qualitative manual annotations. Abbreviations as in Fig 3.

Qualitative results obtained by using the manual annotation tool are highly concordant with quantitative measurements (**Figure 3**). Thus, the qualitative assessment serves as a confirmatory dataset underscoring the robustness of obtained measurement results. Furthermore, it demonstrates the possibility to manually and rapidly tag and categorize tens of thousands of images from complex large-scale datasets using the demonstrated tool, without the necessity of advanced pre-processing and image analysis steps. The manual annotation tool is available on request from *10.5281/zenodo.3367365*, and is compatible with a wide variety of image datasets originating from different imaging modalities.

To further highlight phenotypic diversity and the average morphological response for each compound, the entire dataset was visualized using parallel coordinates plots with each line corresponding to one compound (**Figure 4**). This enables the intuitive detection of trends in the datasets and identification of the number of compounds that deviate from the mean phenotypic change observed in the screen, for both, quantitative morphological measurements (**Figure 4A**) and qualitative manual annotations (**Figure 4B**) in a feature specific and highly comprehensive manner.

In total, about 10% of tested compounds induced an abnormal renal development, affecting either the glomerulus, the tubules or both (**Figures 3, 4**). When D-level ATC groups with a cut-off of a minimum of 4 compounds and ≥ 50% drugs inducing pathological renal phenotypes are chosen, the following compound classes altered pronephric development in zebrafish larvae (**Figure 5, Supplementary Table 1-3**): i) dihydropyridine derivatives (e.g nifedipine (**Figure 5B**)), ii) HMG CoA reductase inhibitors (e.g. atorvastatine (**Figure 5C**)), iii) fibrates (e.g. fenofibrate (**Figure 5D**)), iv) imidazole and triazole derivatives (e.g. sertaconazole (**Figure 5E**)), v) corticosteroids, moderately potent (group II) (e.g. flumethasone (**Figure 5F**)), vi) corticosteroids, potent (group III) (e.g. fluocinolone acetonide (**Figure 5G**)), vii) imidazole derivatives (e.g isoconazole (**Figure 5H**)), viii) glucocorticoids (e.g. methylprednisolone (**Figure 5I**)), ix) acetic acid derivatives and related substances (e.g. diclofenac (**Figure 5J**)), x) propionic acid derivatives (e.g. ibuprofen (**Figure 5K**)), xi) benzimidazole derivatives (e.g. mebendazole (**Figure 5L**)) (see also **Supplementary Tables 1-3**). In addition, several other compounds that did not fulfill above listed ATC group criteria showed notable effects on renal development, such as: proscillaridin A, amiodarone hydrochloride, suloctidil, diltiazem hydrochloride, lidoflazine, irbesartane, ciclopirox ethanolamine, isotretinoin, norgestimate, progesterone, danazol, fludrocortisone acetate, nalidixic acid sodium salt, amphotericin B, miconazole, carmofur, leflunomide, phenylbutazone, piroxicam, meloxicam, mefenamic acid, etofenamate, felbinac, diflunisal, fluspirilen, pimozide, isocarboxazid, disulfiram, nocodoazole, GBR 12909 dihydrochloride, mevastatin, cycloheximide, methiazole, clonixin lysinate, ethoxzolamide, benoxiquine, halofantrine hydrochloride, flunixin meglumine, salmeterol and several others (**Supplementary Table 4**). Obviously, some of the aforementioned compounds chemically belonged to either benzimidazole and imidazole derivates, HMG CoA reductase inhibitors or to non-steroidal anti-inflammatory drug (NSAID) groups and were simply not assigned to an ATC code in the utilized database.

**Figure 5.**
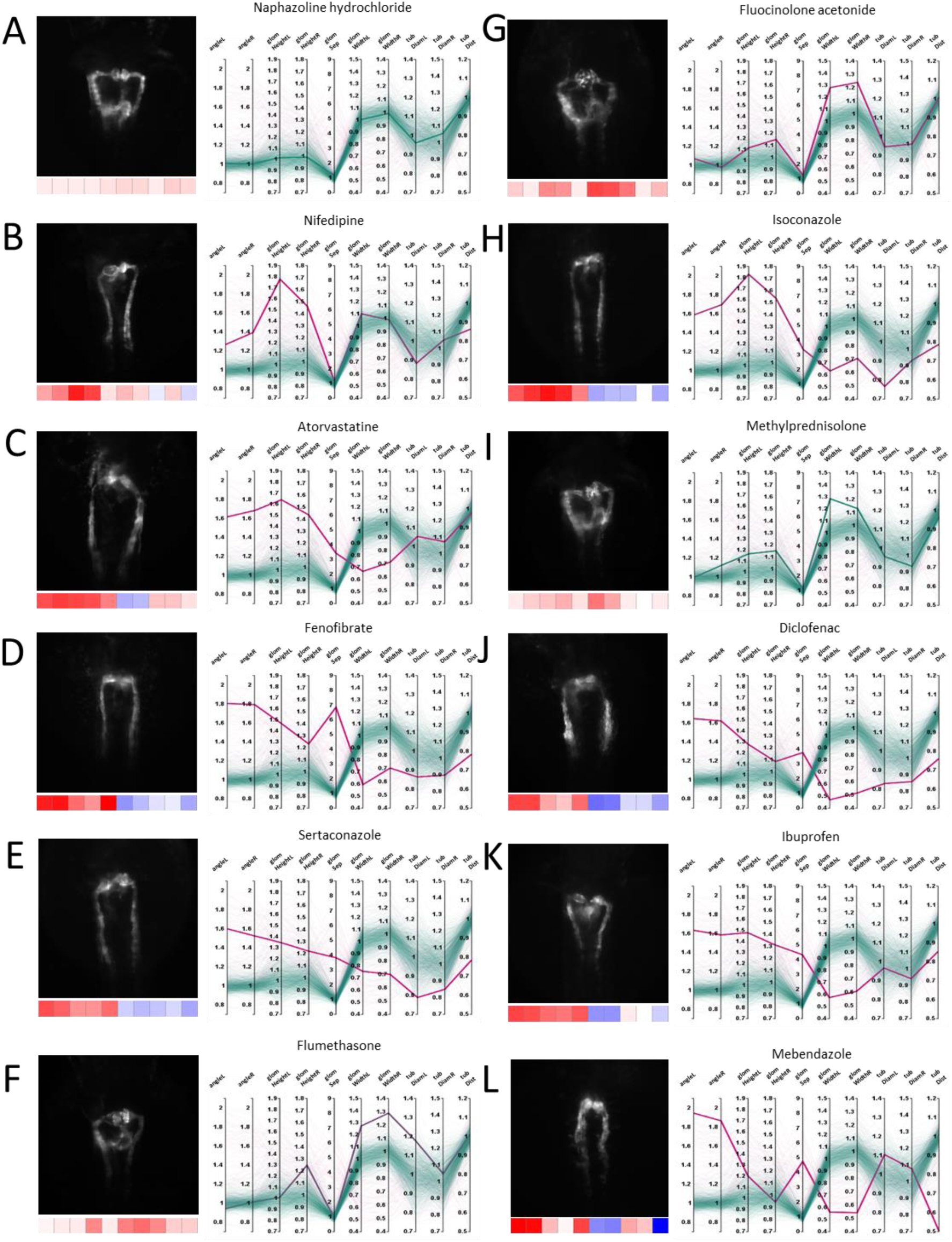
Examples of pronephric phenotypes. Illustrative examples of drug induced phenotypic changes for several compound classes. For each compound a thumbnail image, a heat map z-score visualization (below thumbnail) and a parallel coordinates plot of fold changes of quantitative morphological features are shown. In the parallel coordinates plots, thick lines indicate the shown compound and thin lines represent all other compound treatments; color codes as in Figure 3 and 4. (**A**) Naphazoline hydrochloride (class, sympathomimetics); showing no effect, (**B**) nifedipine (class, dihydropyridine derivatives), (**C**) atorvastatine (class, HMG CoA reductase inhibitors), (**D**) fenofibrate (class, fibrates), (**E**) sertaconazole (class, imidazole and triazole derivatives), (**F**) flumethasone (class, corticosteroids, moderately potent (group II)), (**G**) fluocinolone acetonide (class, corticosteroids, potent (group III)), (**H**) isoconazole (class, imidazole derivatives), (**I**) Methylprednisolone (class, Glucocorticoids), (**J**) Diclofenac (class, Acetic acid derivatives and related substances), (**K**) Ibuprofen (class, Propionic acid derivatives), and (L) Mebendazole (class, Benzimidazole derivatives).

The detailed list of results is shown in **Supplementary Table 1**. To facilitate the further analysis of the dataset and provide a resource for investigation also by other researchers, we have generated an interactive exploration tool that allows intuitively browsing and searching the entire dataset (**Figure 6** and **Supplementary Software 4**). The tool consists of a scatter plot of a 3-dimensional PCA of normalized quantitative features (**Figure 6A**), a parallel coordinates plot of fold changes based on quantitative measurements (**Figure 6B**), sliders to limit the displayed data based on 3D principal component analysis (PCA) results (not shown) and the full results table (**Figure 6C**). All elements are interactive and allow highlighting single compounds or compound classes by selecting plot entries, moving of sliders or by searching the result table, as illustrated in **Figure 6** for the class of propionic acid derivatives (ATC code D-Level M01AE). The exploration tool can be launched by simply executing the provided KNIME workflow with the provided data table (**Figure 6D** and see **Supplementary Software 4**). Ultimately, the dataset can also be visualized in the tensorflow-projector web interface (https://projector.tensorflow.org/), which also offers interactive browsing functionality using a 3D scatter plot derived from dimensionality reduction of the morphological feature dataset. Additional methods for dimensionality reduction other than PCA are available, namely UMAP and T-SNE, which are not evaluated here. An advantage of the tensorflow projector interface over the KNIME exploration workflow, is the built-in nearest-neighbor search which allows to identify compounds closest to a query compound in term of morphological features profile. This functionality is illustrated in **Supplementary Figure 2** with S-(+)-ibuprofen as a query compound.

**Figure 6.**
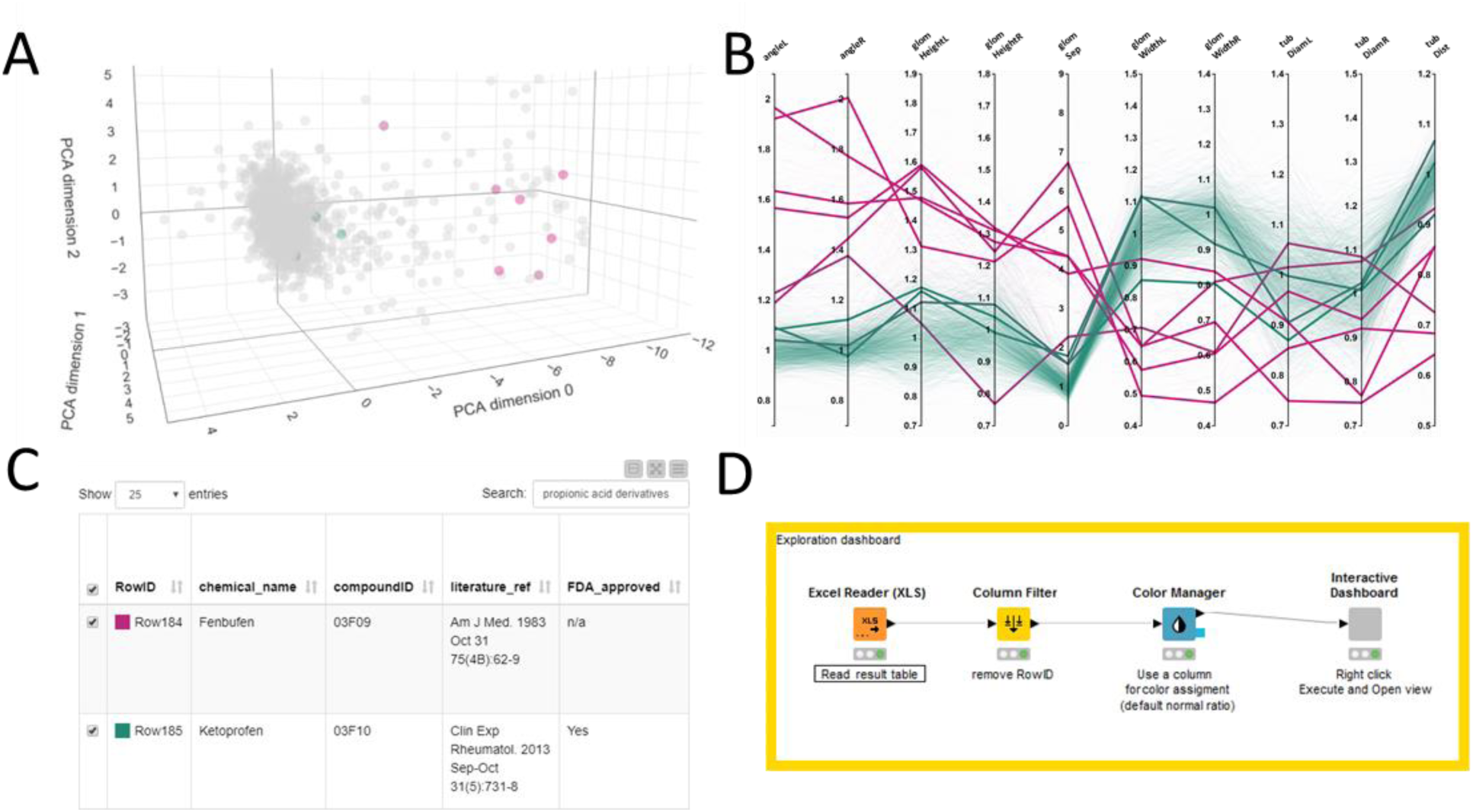
Exploring and browsing the generated zebrafish embryo nephrotoxicity dataset. Shown are screenshots of an interactive exploration tool generated using KNIME (see Supplementary Software 4). The workflow allows to visualize the full result table as well as a PCA scatter plot and a parallel coordinates line plot of the quantitative features in a single HTML window. The tool allows highlighting single compounds, compound groups or classes by selecting single lines in a parallel coordinates plot, sliders for limiting PCA range (not shown) or through searching in the data result table. Selected results are shown as colored dots in the PCA scatterplot, thicker lines in the parallel coordinates plot or as filtered rows in the results table. In this figure all panels show highlighted results for the propionic acid derivatives compound class. (**A**) PCA plot, (**B**) parallel coordinate plot, (**C**) result table. Color code for highlighted dots, lines or rows in A-C as in Figures 3 and 4 with 0% (magenta, all abnormal) to 100% (green, all normal). (**D**) KNIME workflow generating the interactive tool.

### General phenotyping

Brightfield images of zebrafish larvae were taken following drug-exposure to assess extrarenal morphological anomalies and phenotypic indicators of general toxicity. These parameters included number/percentage of dead larvae, larvae having a curved back or tail, and larvae showing either mild or severe edema. Further non-renal parameters included the presence of yolk sac necrosis, malformed somites, and heartbeat alterations (faster, slower, absent). In addition, from fluorescence images, absence of a pancreatic signal was recorded (**Supplementary Table 1**).

For instance, imidazoles, triazole derivatives and benzimidazole derivatives provoked high degrees of upward-curved tails in larval zebrafish. Especially in imidazole and triazole derivatives, this was accompanied by mild pericardial edema. Dihydropyridine derivatives (selective calcium channel blockers) and HMG CoA reductase inhibitors mainly provoked pericardial edema. Of note, there was no strong correlation between the development of pericardial edema and the presence of renal alterations (**Supplementary Figure 1**).

## Discussion

Drug safety is of utmost importance, especially in pregnancy and in very premature neonates with ongoing organogenesis. Toxic exposure may potentially result in impaired development with life-long consequences. However, knowledge on drug-induced developmental toxicology is very limited for a multitude of approved drugs, and safety information in human is often largely based on observational data. For new drugs, mandatory embryo-fetal developmental and reproductive toxicity (DART) studies for safety assessment are generally restricted to rat and rabbits (Ishihara-Hattori and Barrow, 2016). Large numbers of animals are required for these studies; thus, they are complex, cost-intensive, and time-consuming. Hence, there is an unmet need for large-scale *in vivo* developmental toxicity drug screening investigations in appropriate animal models. In the context of the three R’s, current legislation promotes the use of alternative animal models including zebrafish embryos.

In this study, we therefore expanded on our previously published protocol for automated acquisition of bilateral symmetric organs in zebrafish embryos and robustly scored the morphological alterations of the pronephros in a large-scale zebrafish chemical screen (Pandey et al., 2019, Westhoff et al., 2013). Consistent dorsal positioning of embryos was achieved through utilization of 3D-printed orientation tools (Wittbrodt et al., 2014), and microplates containing arrayed embryos. Samples were captured using an automated widefield screening microscope equipped with stationary sample holder and moving optics to avoid any change of orientation of specimen during automated image acquisition (Pandey et al., 2019). Using this pipeline, we were able to efficiently capture consistent dorsal views of embryonic kidneys of more than 15,000 embryos treated with a library of 1,280 off-patent (> 95% approved) drugs within 8 months, with on average one imaging day per week. Imaging itself takes only about 12 min per 96-well plate. Limitations restricting higher throughput are labor-intensive manual embryo generation, handling and drug treatment procedures. While advanced automated drug screening technologies exist, full robotic automation of compound or plate handling is often economically unjustified in zebrafish screening routines. Manual orientation within agarose filled microplates takes 15-20 min per 96 embryos causing a significant workload; however, one plate can be prepared while another plate is automatically imaged, so this has not significantly impaired overall throughput. Nevertheless, technical solutions for fully automated orientation and imaging of zebrafish embryos exist, but seem to lack the throughput required to image several hundred embryos within a few hours (Early et al., 2018).

Using automated image pre-processing and semi-automated analysis tools, this highly standardized image datasets (**Figure 1B**) allowed the qualitative categorization of renal phenotypes and the quantitative morphological analysis based on simple geometric measurements. This led to a numerical fingerprint (**Figure 3, Figure4**, and **Figure 5**) for each tested substance enabling the scoring of slight and gross morphological alterations of the developing pronephros upon drug treatment. Each embryo data were analyzed by a single expert operator using the above tools, causing a major analysis workload but also generating a fully annotated and highly gauged high-content screening dataset. Undoubtedly, it would be highly beneficial to develop automated image analysis solutions that allow morphological differentiation of pronephric phenotypes. However, developing algorithms for automated and robust scoring of complex phenotypes in biologically heterogeneous whole-organism screening datasets is a highly challenging task. We are currently exploring the usage of machine- and deep-learning methodologies taking advantage of this work as a benchmark and validation dataset.

Results from our screen revealed a set of compound classes that provoked renal developmental toxicity. Major substance classes inducing severe pronephric malformations included dihydropyridine derivatives (calcium channel blockers), HMG CoA reductase inhibitors, fibrates, imidazole and triazole derivatives (e.g. antifungals for topical use), moderately potent and potent corticosteroids, glucocorticoids, acetic acid derivatives and related substances, propionic acid derivatives, and benzimidazole derivatives (i.e. antinematodal agents) (**Supplementary Table 3**). Single compounds that in the utilized version of the ATC code are not assigned to a D-level also revealed notable detrimental effects on renal development were also identified (**Supplementary Table 4**). A large fraction of the drugs identified in our screen recapitulate human observational or experimental data including not only renal but overall developmental toxicity, embryotoxicity, fetotoxicity, and/or teratogenicity.

Several compounds from the dihydropyridine derivatives class, one out of three chemical groups of calcium channel blockers, revealed renal developmental toxicity in zebrafish. For example, the most prescribed calcium channel blocker nifedipine caused strong tubular and glomerular malformations (**Figure 1C**). Although not licensed, nifedipine is regularly used in clinical practice for the tocolytic management of preterm labor and for pregnancy-induced hypertensive disorders. To our knowledge, so far there is no detailed published data on renal developmental toxicity for dihydropyridine derivatives. However, there is mixed observational human data on *in-utero* exposure to calcium channel blockers and increased perinatal mortality, increased odds of preterm birth and perinatal mortality, and evidence of increased malformations, including hypospadias (Fitton et al., 2017).

HMG CoA reductase inhibitors showed strong detrimental effects on pronephros development. They are not recommended in early pregnancy. While cholesterol is known to be essential for fetal development, a systematic review showed no clear relationship with statin use and congenital anomalies in pregnancy (Karalis et al., 2016); however, data was inconclusive.

Fibrates prominently displayed morphological pronephros alterations. In humans, they are known to cross the human blood– placenta barrier (Tsai et al., 2004). Animal reproduction studies demonstrated adverse effects on the fetus, however, no teratogenicity was noted in several case reports of fibrate use after the first trimester (Wong et al., 2015). While fibrate-associated nephrotoxicity has been described in adults, to our knowledge there is no data on fetal developmental nephrotoxicity (Attridge et al., 2013).

Corticoid drugs, widely prescribed e.g. to very preterm neonates, showed very subtle but consistent phenotypes in our screen. Excessive glucocorticoid signaling is detrimental for fetal development; slowing fetal and placental growth and programming the individual for disease later in adult life (Busada and Cidlowski, 2017). In addition, first-trimester corticosteroid exposure, including dermatologic steroids, slightly increases the risk of orofacial clefts (Xiao et al., 2017). Interestingly, a nephron deficit following elevated maternal glucocorticoid exposure and leading to hypertension in adult animals was observed in fetal rats and sheep (Moritz et al., 2011, Ortiz et al., 2003, Singh et al., 2007).

Azoles were especially noticeable in our screen. They are known to produce reproductive and developmental toxicity in both human and animals. Azole fungicides, classified into triazoles and imidazoles, inhibit key enzymes in steroidogenesis that, among others, provoke embryotoxicity and teratogenic malformations (Zarn et al., 2003). In rats, antifungal azoles were teratogenic producing fetal defects such as cleft palate and hydronephrosis/hydroureter (Pursley et al., 1996). Various benzimidazole antihelmintics, including mebendazole, albendazole and flubendazole have, to different degrees though, been associated with embryofetal developmental toxicity *in vivo* in different animals including zebrafish, and *in vitro* (Carlsson et al., 2011, Dayan, 2003, Longo et al., 2013, Sasagawa et al., 2016). Information on renal developmental toxicity, however, is scarce.

Most nonsteroidal anti-inflammatory drugs (NSAIDs) assayed in our screen caused severe renal malformations in zebrafish embryo. NSAIDS are often taken in pregnancy for acute pain or chronic conditions such as rheumatologic disorders, but are also administered to preterm neonates for closure of a patent ductus arteriosus. Early NSAID exposure has been reported to increase spontaneous abortion rate and possibly cause congenital malformations (Nielsen et al., 2001). NSAID-induced renal effects can in rare instances be severe, particularly after 32 weeks’ gestation, with potential for neonatal renal failure (Morgan et al., 2014). However, the magnitude of risk in various clinical scenarios remains unclear and may vary with the dose, duration, timing of therapy, and maternal indication for use. Of note, in zebrafish pronephroi, it has been demonstrated that prostaglandin signaling, i.e. PGE2 signaling, regulates nephron formation including proximal and distal segment formation during nephrogenesis (Chambers and Wingert, 2019).

From the list of not ATC D-level assigned drugs that provoked developmental nephrotoxicity in our screen, several have been associated with developmental and/or adult nephrotoxicity in previous investigations, in particular amphotericin B (Hanna et al., 2016), phenylbutazone (Brix, 2002) and piroxicam and meloxicam (Boubred et al., 2006). Of note, several drugs (e.g. diltiazem hydrochloride, lidoflazine, etofenamate, felbinac, mevastatin, methiazole, flunixin meglumine) could also be assigned to ATC D-level groups that were previously shown to induce renal abnormalities in our screening investigation. Irbesartane is a member of angiotensin II receptor type 1 (AT1-R) antagonists. For this group of drugs as well as for ACE inhibitors, experimental data and human observations argue for developmental nephrotoxicity (Boubred et al., 2006). In our screening experiment, however, in contrast to other relevant groups only minor alterations of renal development were observed. The reason for this will be speculated on in the following paragraph.

There are several limitations to our study. First, considering the workload of this project, we had to choose one single drug concentration for all compounds. In fact, concentration series for drug classes with only minor effects (such as ACE inhibitors and AT1-R antagonists) might unravel stronger toxic effects using different concentrations. As such, we have previously shown that captopril and losartan provoked developmental nephrotoxicity with increasing concentration (Westhoff et al., 2013). Hence, detailed secondary analyses of the groups identified in our screen are currently being carried out in our laboratory. Second, the zebrafish embryos were drug-exposed via the fish water. The transdermal uptake of compounds into the zebrafish can vary depending on the chemical properties of the specific drug. This has already been shown e.g. for gentamicin, a member of aminoglycoside antibiotics, that demands very high concentrations when applied via fish water exposure in contrast to microinjection (Gorgulho et al., 2018, Hentschel et al., 2005, Westhoff et al., 2013). Additionally, while there are substantial structural and physiological similarities between nephrons of the zebrafish pronephros and the mammalian metanephros, the timing of nephron differentiation and kidney development as well as associated spatio-temporal gene-expression patterns differ between the zebrafish and other vertebrates (Gerlach and Wingert, 2013). This must be taken into account when considering the very short time period for pronephros development in zebrafish and the proportionally long time period of drug exposure in our study.

## Conclusion

In conclusion, the results of our developmental nephrotoxicity screen reconfirm previous data from animal experiments and human observational data regarding a variety of drugs, but also put additional substance classes into focus. Thus, our data contributes to the knowledge on approved drugs with organo-toxic side-effects during vertebrate organogenesis that was previously unavailable. In combination with the generated data exploration tool (**Figure 6**), the data provides a compelling resource with high information content to other bio-medical researchers for further exploration or as a starting point of follow-up studies. Moreover, it might serve as a catalog for health professional to identify substances of potential concern when used by pregnant women or in preterms. This, in the end, might have an impact on the prescription of drugs during pregnancy and on the drug-exposition of very preterm neonates. Since several renal congenital anomalies, i.e. renal hypoplasia and dysplasia, are at least partially assigned to unfavorable fetal environmental exposures including nephrotoxic chemical compounds, the identified drugs of our screen require further investigation in the future. Ultimately, the demonstrated workflow and associated tools are not restricted to the analysis of the embryonic kidney and can be readily adapted to conduct other organ- or tissue-specific large-scale chemical screening experiments in the zebrafish embryo model.

## Materials and Methods

### Ethics statement

The work presented does not involve work with animals according to German and European legislation. To obtain zebrafish embryos and larvae, fish were maintained in closed stocks at the Karlsruhe Institute of Technology (KIT). All zebrafish husbandry and experimental procedures were performed in accordance with the German animal protection regulations (Regierungspräsidium Karlsruhe, Germany; Tierschutzgesetz 111, Abs. 1, Nr. 1, AZ35-9185.64/BH). The facility is under the supervision of the Regierungspräsidium Karlsruhe. The *Tg(wt1b:EGFP)* transgenic line has been previously described (Perner et al., 2007).

### Fish keeping, embryo handling and drug exposure

Adult zebrafish of the *Tg(wt1b:EGFP)* transgenic line (Perner et al., 2007) were kept and maintained according to standard protocols (Westerfield, 2000a). Eggs were collected from pairwise and batch crossings in cages equipped with dividers that were removed before eggs were required (Westerfield, 2000b). The developmental stage of embryos was determined as previously described (Kimmel et al., 1995). Embryos were raised in fish water at 28.5°C in E3 medium. At 24 hours post fertilization (hpf), green fluorescent protein (GFP) positive embryos were collected and enzymatically dechorionated using 10 mg/mL Pronase (Sigma Aldrich, Taufkirchen, Germany) (Gehrig et al., 2009). Embryos were transferred to a beaker, washed 3 times with 400 mL of fish water and transferred to 12-well plates containing 2 mL of a 25 µM compound solution of the Prestwick Chemical Library® (Prestwick Chemical, D’Illkirch, France) in 0.5% DMSO in 5 mM HEPES-buffered E3 medium supplemented with 0.003% 1-pheny-2-thiourea (PTU, Alfa Aesar, Karlsruhe, Germany). This library contains 1,280 off-patent small molecules, 95% approved drugs (FDA, EMA and other agencies), dissolved in DMSO. Following a 24h drug exposure at 28.5°C, the solutions were removed at 48 hpf, embryos were washed and kept in E3/PTU containing 250 µg/mL tricaine. To score overall morphological phenotypes, brightfield overview images were taken using a stereo microscope after ending of compound exposure. Subsequently, larvae were transferred to agarose-filled 96-well plates for kidney imaging. Embryos treated with 0.5% DMSO served as negative controls.

### Preparation of agarose molds in microtiter plates

48 hpf embryos were oriented in agarose-filled 96-well plates generated with 3D-printed orientation tools as previously described (Pandey et al., 2019, Wittbrodt et al., 2014). In brief, each well of a 96-well microtiter plate (Cat. No. 655101, Greiner, Frickenhausen, Germany) was filled with 60 µL of 1% agarose in E3 medium supplemented with 250 µg/mL tricaine (Sigma Aldrich, Taufkirchen, Germany) using a multi-channel pipette. Cavities for dorsal orientation were generated by inserting a 3D-printed orientation tool (Wittbrodt et al., 2014). The embryos were oriented dorsally under a stereomicroscope.

### Image acquisition

96-well microtiter plates containing zebrafish embryos were automatically imaged on an ACQUIFER Imaging Machine (ACQUIFER Imaging GmbH, Heidelberg, Germany) widefield high-content screening microscope equipped with a white LED (light-emitting diode) array for brightfield imaging, a LED fluorescence excitation light source, a sCMOS (2048 × 2048 pixel) camera, a stationary plate holder in combination with moving optics and a temperature-controlled incubation lid. Pronephric areas were imaged in the brightfield and 470 nm channels using 10 z-slices (dZ = 15 µm) and a 4x NA 0.13 objective (Nikon, Düsseldorf, Germany). The focal plane to center the z-stack was detected in the 470 nm channel using a built-in software autofocus algorithm. Integration times were fixed at 30% relative LED intensity and 20 ms exposure time for the brightfield channel and 100% relative LED intensity and 100 ms exposure time for the 470 nm channel. Imaging times were approximately 12 min for a full 96-well plate.

### Data handling and visualization

Image data was stored and processed on an ACQUIFER HIVE (ACQUIFER Imaging GmbH, Heidelberg, Germany). Raw images of fluorescence channel were processed using a custom written Perl script in combination with a Fiji macro (**Supplementary Software 1** and **2**). This script and Fiji macro generated multi-layer z-stacks, XY-cropped maximum projections for each experimental embryo and thumbnail montage images for each experimental microplate readily allowing visual assessment and comparison of pronephric phenotypes as described in (Pandey et al., 2019, Westhoff et al., 2013). In total, 4.2 TB of data corresponding to >15,000 embryos treated with 1,280 compounds were acquired and processed.

### Scoring of pronephric phenotypes

To score phenotypic alterations upon compound treatment, 26 parameters were scored for each compound treatment. Extrarenal parameters included mortality, curved back/tail, mild and severe edema, heartbeat alterations, somite malformations, and yolk sac necrosis and were assigned based on manual evaluation of treated embryos on a stereo microscope prior to mounting in agarose-filled microplates. Screening datasets were evaluated using qualitative manual scoring of kidney phenotypes by assignment of up to 10 different phenotypic categories: reduced pronephros angle major (rpa_maj), reduced pronephros angle moderate (rpa_mod), reduced pronephros angle minor (rpa_min), no fluorescence (empty), glomerular malformation (glom_malform), glomerular separation major (glomsep_maj), glomerular separation moderate (glomsep_mod), glomerular separation minor (glomsep_min), impaired liver pancreas area (liver-panc_pheno) and normal kidney (normal_kidney). This assignment was carried out using a manual annotation tool (MAT) developed at ACQUIFER (10.5281/zenodo.3367365). The MAT tool allows the rapid manual assignment (blind or non-blind) of user defined categories to screening datasets after user-defined image data dimensionality reduction (z-projections, auto-cropping) and visualization improvements (look-up tables, intensity scaling). In summary, each well coordinate was assigned one or multiple categories corresponding to qualitative descriptions of phenotypic alterations of the kidney. Quantitative measurements of kidney alterations were carried out for 10 morphological pronephric parameters: pronephric angle left (angleL), pronephric angle right (angleR), glomerular height left (glomHeightL), glomerular height right (glomHeightR), glomerular separation (glomSep), glomerular width left (glomWidthL), glomerular width right (glomWidthR), tubular diameter left (tubDiamL), tubular diameter right (tubDiamR) and tubular distance (tubDist). To measure these features, cropped maximum projection data of fluorescently labeled kidneys were loaded in Fiji and 16 reference points were manually set and the geometrical parameters (distances and angles) were automatically calculated using a Fiji macro (**Figure 2** and **Supplementary Software 1**).

### Data analysis

For all data analysis steps KNIME (Mazanetz et al., 2012) was used and workflows are available in (**Supplementary Software 3**). Result files from gross morphological scoring, qualitative manual annotation and quantitative phenotypic measurements were processed and combined with information about the Prestwick library and Anatomical Therapeutic Chemical (ATC) Classification System information. Data for the Prestwick library was received from Prestwick Chemical, d’Illkirch, France, and ATC classification data was downloaded from KEGG: Kyoto Encyclopedia of Genes and Genomes database (Kanehisa and Goto, 2000). In brief, the workflow loads the csv files containing measurement data from each of the 199 experimental folders. To remove extreme outliers caused by damaged, misaligned or severely malformed embryos, data points more than 2 standard deviations (SD) away from the mean of a specific measurement were excluded from further analysis. To minimize experimental variations, each quantitative measurement value was divided by the mean of negative controls from the same experimental day, leading to feature specific fold change values. To further facilitate cross-compound comparison, fold changes were z-score normalized. For qualitative phenotypic categories and gross morphological phenotypes, the sum of embryos per experimental treatment assigned with a certain category was calculated. The full dataset was saved as XLS-file (**Supplementary Tables 1,2**). Heat map visualizations were generated using matrix2png (Pavlidis and Noble, 2003). Principal component analysis (PCA) was used for dimensionality reduction from the 10 normalized morphological features down to 3 dimensions for visualization as a 3D scatter plot. Using the first 3 PCA components results in an approximation of the initial datasets representing 76% of the initial data variance. A PCA scatter plot, a parallel coordinates plot and original data table were rendered as a single interactive dashboard to enable convenient data browsing and visualization: selecting rows in the data table triggers highlighting in the parallel coordinates and scatter plot, and *vice versa*. The interactive dashboard is generated in KNIME (**Supplementary Software 4**). The dataset was also visualized as a 3D PCA scatter plot on the online tensorflow-projector platform (https://projector.tensorflow.org/) (**Supplementary Figure 2** and **Supplementary Information**), which offers additional projection methods (UMAP, T-SNE) and functions to identify the closest compounds from a selected compound.

## Supporting information

Supplementary Software

## Acknowledgments

We are grateful to Felix Loosli (KIT) and Franz Schaefer (Children’s Hospital Heidelberg) for general support, and Dr Christoph Englert (Leibniz Institute for Age Research-Fritz Lipmann Institute, Jena, Germany) for kindly providing the *Tg(wt1b:EGFP)* transgenic line. We thank Jonas Wagner (KIT) for python programming. The authors acknowledge the excellent fish husbandry of Nadeshda Wolf and Natalja Kusminski.

## Author contributions

Conceived and designed the experiments: JHW, PJS, JG. Performed the experiments: PJS, LC. Analyzed the data: JHW, PJS, LSVT, JH, TB, JG. Wrote the manuscript: JHW, LSVT, JH, JG. Generated data handling, image analysis and visualization tools: PJS, LSVT, JG. Designed the study: JHW, JG. Gave general advice and edited the manuscript: GFH, BT. Supervised the study: JHW, JG.

## References

Attridge, R. L., Frei, C. R., Ryan, L., Koeller, J. & Linn, W. D. 2013. Fenofibrate-associated nephrotoxicity: a review of current evidence. Am J Health Syst Pharm, 70, 1219-25. 10.2146/ajhp120131

Ball, J. S., Stedman, D. B., Hillegass, J. M., Zhang, C. X., Panzica-Kelly, J., Coburn, A., Enright, B. P., Tornesi, B., Amouzadeh, H. R., Hetheridge, M., Gustafson, A. L. & Augustine-Rauch, K. A. 2014. Fishing for teratogens: a consortium effort for a harmonized zebrafish developmental toxicology assay. Toxicol Sci, 139, 210-9. 10.1093/toxsci/kfu017

Boubred, F., Vendemmia, M., Garcia-Meric, P., Buffat, C., Millet, V. & Simeoni, U. 2006. Effects of maternally administered drugs on the fetal and neonatal kidney. Drug Saf, 29, 397-419. 10.2165/00002018-200629050-00004

Brady, C. A., Rennekamp, A. J. & Peterson, R. T. 2016. Chemical Screening in Zebrafish. Methods Mol Biol, 1451, 3-16. 10.1007/978-1-4939-3771-4_1

Brannen, K. C., Panzica-Kelly, J. M., Danberry, T. L. & Augustine-Rauch, K. A. 2010. Development of a zebrafish embryo teratogenicity assay and quantitative prediction model. Birth Defects Res B Dev Reprod Toxicol, 89, 66-77. 10.1002/bdrb.20223

Brent, R. L. 2004. Environmental causes of human congenital malformations: the pediatrician’s role in dealing with these complex clinical problems caused by a multiplicity of environmental and genetic factors. Pediatrics, 113, 957–68.

Brix, A. E. 2002. Renal papillary necrosis. Toxicol Pathol, 30, 672-4. 10.1080/01926230290166760

Busada, J. T. & Cidlowski, J. A. 2017. Mechanisms of Glucocorticoid Action During Development. Curr Top Dev Biol, 125, 147-170. 10.1016/bs.ctdb.2016.12.004

Carlsson, G., Patring, J., Ulleras, E. & Oskarsson, A. 2011. Developmental toxicity of albendazole and its three main metabolites in zebrafish embryos. Reprod Toxicol, 32, 129-37. 10.1016/j.reprotox.2011.05.015

Chambers, B. E. & Wingert, R. A. 2019. Mechanisms of Nephrogenesis Revealed by Zebrafish Chemical Screen: Prostaglandin Signaling Modulates Nephron Progenitor Fate. Nephron, 143, 68-76. 10.1159/000501037

Dayan, A. D. 2003. Albendazole, mebendazole and praziquantel. Review of non-clinical toxicity and pharmacokinetics. Acta Trop, 86, 141–59.

De Vigan, C., De Walle, H. E., Cordier, S., Goujard, J., Knill-Jones, R., Ayme, S., Calzolari, E. & Bianchi, F. 1999. Therapeutic drug use during pregnancy: a comparison in four European countries. OECM Working Group. Occupational Exposures and Congenital Anomalies. J Clin Epidemiol, 52, 977-82. S0895435699000918 [pii]

Drummond, I. A. 2005. Kidney development and disease in the zebrafish. J Am Soc Nephrol, 16, 299-304. ASN.2004090754 [pii] 10.1681/ASN.2004090754

Drummond, I. A. & Davidson, A. J. 2010. Zebrafish kidney development. Methods Cell Biol, 100, 233-60. B978-0-12-384892-5.00009-8 [pii] 10.1016/B978-0-12-384892-5.00009-8

Early, J. J., Cole, K. L., Williamson, J. M., Swire, M., Kamadurai, H., Muskavitch, M. & Lyons, D. A. 2018. An automated high-resolution in vivo screen in zebrafish to identify chemical regulators of myelination. Elife, 7. 10.7554/eLife.35136

Fitton, C. A., Steiner, M. F. C., Aucott, L., Pell, J. P., Mackay, D. F., Fleming, M. & Mclay, J. S. 2017. In-utero exposure to antihypertensive medication and neonatal and child health outcomes: a systematic review. J Hypertens, 35, 2123-2137. 10.1097/HJH.0000000000001456

Gehrig, J., Reischl, M., Kalmar, E., Ferg, M., Hadzhiev, Y., Zaucker, A., Song, C., Schindler, S., Liebel, U. & Muller, F. 2009. Automated high-throughput mapping of promoter-enhancer interactions in zebrafish embryos. Nat Methods, 6, 911-6. nmeth.1396 [pii] 10.1038/nmeth.1396

Gerlach, G. F. & Wingert, R. A. 2013. Kidney organogenesis in the zebrafish: insights into vertebrate nephrogenesis and regeneration. Wiley Interdiscip Rev Dev Biol, 2, 559-85. 10.1002/wdev.92

Gorgulho, R., Jacinto, R., Lopes, S. S., Pereira, S. A., Tranfield, E. M., Martins, G. G., Gualda, E. J., Derks, R. J. E., Correia, A. C., Steenvoorden, E., Pintado, P., Mayboroda, O. A., Monteiro, E. C. & Morello, J. 2018. Usefulness of zebrafish larvae to evaluate drug-induced functional and morphological renal tubular alterations. Arch Toxicol, 92, 411-423 10.1007/s00204-017-2063-1

Gunnarsson, L., Jauhiainen, A., Kristiansson, E., Nerman, O. & Larsson, D. G. 2008. Evolutionary conservation of human drug targets in organisms used for environmental risk assessments. Environ Sci Technol, 42, 5807–13.

Hanna, M. H., Askenazi, D. J. & Selewski, D. T. 2016. Drug-induced acute kidney injury in neonates. Curr Opin Pediatr, 28, 180-7. 10.1097/MOP.0000000000000311

Hentschel, D. M., Park, K. M., Cilenti, L., Zervos, A. S., Drummond, I. & Bonventre, J. V. 2005. Acute renal failure in zebrafish: a novel system to study a complex disease. Am J Physiol Renal Physiol, 288, F923-9. 00386.2004 [pii] 10.1152/ajprenal.00386.2004

Hinsberger, A., Wingen, A. M. & Hoyer, P. F. 2001. Angiotensin-II-receptor inhibitors in pregnancy. Lancet, 357, 1620.

Howe, K., Clark, M. D., Torroja, C. F., Torrance, J., Berthelot, C., Muffato, M., Collins, J. E., Humphray, S., Mclaren, K., Matthews, L., Mclaren, S., Sealy, I., Caccamo, M., Churcher, C., Scott, C., Barrett, J. C., Koch, R., Rauch, G. J., White, S., Chow, W., Kilian, B., Quintais, L. T., Guerra-Assuncao, J. A., Zhou, Y., Gu, Y., Yen, J., Vogel, J. H., Eyre, T., Redmond, S., Banerjee, R., Chi, J., Fu, B., Langley, E., Maguire, S. F., Laird, G. K., Lloyd, D., Kenyon, E., Donaldson, S., Sehra, H., Almeida-King, J., Loveland, J., Trevanion, S., Jones, M., Quail, M., Willey, D., Hunt, A., Burton, J., Sims, S., Mclay, K., Plumb, B., Davis, J., Clee, C., Oliver, K., Clark, R., Riddle, C., Elliot, D., Threadgold, G., Harden, G., Ware, D., Begum, S., Mortimore, B., Kerry, G., Heath, P., Phillimore, B., Tracey, A., Corby, N., Dunn, M., Johnson, C., Wood, J., Clark, S., Pelan, S., Griffiths, G., Smith, M., Glithero, R., Howden, P., Barker, N., Lloyd, C., Stevens, C., Harley, J., Holt, K., Panagiotidis, G., Lovell, J., Beasley, H., Henderson, C., Gordon, D., Auger, K., Wright, D., Collins, J., Raisen, C., Dyer, L., Leung, K., Robertson, L., Ambridge, K., Leongamornlert, D., Mcguire, S., Gilderthorp, R., Griffiths, C., Manthravadi, D., Nichol, S., Barker, G., et al. 2013. The zebrafish reference genome sequence and its relationship to the human genome. Nature, 496, 498-503. 10.1038/nature12111

Ishihara-Hattori, K. & Barrow, P. 2016. Review of embryo-fetal developmental toxicity studies performed for recent FDA-approved pharmaceuticals. Reprod Toxicol, 64, 98-104. 10.1016/j.reprotox.2016.04.018

Kanehisa, M. & Goto, S. 2000. KEGG: kyoto encyclopedia of genes and genomes. Nucleic Acids Res, 28, 27-30 10.1093/nar/28.1.27

Karalis, D. G., Hill, A. N., Clifton, S. & Wild, R. A. 2016. The risks of statin use in pregnancy: A systematic review. J Clin Lipidol, 10, 1081-90. 10.1016/j.jacl.2016.07.002

Kimmel, C. B., Ballard, W. W., Kimmel, S. R., Ullmann, B. & Schilling, T. F. 1995. Stages of embryonic development of the zebrafish. Dev Dyn, 203, 253-310. 10.1002/aja.1002030302

Kirk, R. G. W. 2018. Recovering The Principles of Humane Experimental Technique: The 3Rs and the Human Essence of Animal Research. Sci Technol Human Values, 43, 622-648 10.1177/0162243917726579

Longo, M., Zanoncelli, S., Colombo, P. A., Harhay, M. O., Scandale, I., Mackenzie, C., Geary, T., Madrill, N. & Mazue, G. 2013. Effects of the benzimidazole anthelmintic drug flubendazole on rat embryos in vitro. Reprod Toxicol, 36, 78-87. 10.1016/j.reprotox.2012.12.004

Mackenzie, H. S. & Brenner, B. M. 1995. Fewer nephrons at birth: a missing link in the etiology of essential hypertension? Am J Kidney Dis, 26, 91–8.

Macrae, C. A. & Peterson, R. T. 2015. Zebrafish as tools for drug discovery. Nat Rev Drug Discov, 14, 721-31 10.1038/nrd4627

Mastrobattista, J. M. 1997. Angiotensin converting enzyme inhibitors in pregnancy. Semin Perinatol, 21, 124–34.

Mazanetz, M. P., Marmon, R. J., Reisser, C. B. & Morao, I. 2012. Drug discovery applications for KNIME: an open source data mining platform. Curr Top Med Chem, 12, 1965-79. 10.2174/156802612804910331

Morgan, T. M., Jones, D. P. & Cooper, W. O. 2014. Renal teratogens. Clin Perinatol, 41, 619-32. 10.1016/j.clp.2014.05.010

Moritz, K. M., De Matteo, R., Dodic, M., Jefferies, A. J., Arena, D., Wintour, E. M., Probyn, M. E., Bertram, J. F., Singh, R. R., Zanini, S. & Evans, R. G. 2011. Prenatal glucocorticoid exposure in the sheep alters renal development in utero: implications for adult renal function and blood pressure control. Am J Physiol Regul Integr Comp Physiol, 301, R500-9. 10.1152/ajpregu.00818.2010

Nagel, R. 2002. DarT: The embryo test with the Zebrafish Danio rerio--a general model in ecotoxicology and toxicology. ALTEX, 19 Suppl 1, 38–48.

Nielsen, G. L., Sorensen, H. T., Larsen, H. & Pedersen, L. 2001. Risk of adverse birth outcome and miscarriage in pregnant users of non-steroidal anti-inflammatory drugs: population based observational study and case-control study. BMJ, 322, 266-70. 10.1136/bmj.322.7281.266

Ortiz, L. A., Quan, A., Zarzar, F., Weinberg, A. & Baum, M. 2003. Prenatal dexamethasone programs hypertension and renal injury in the rat. Hypertension, 41, 328–34.

Pandey, G., Westhoff, J. H., Schaefer, F. & Gehrig, J. 2019. A Smart Imaging Workflow for Organ-Specific Screening in a Cystic Kidney Zebrafish Disease Model. Int J Mol Sci, 20. 10.3390/ijms20061290

Pavlidis, P. & Noble, W. S. 2003. Matrix2png: a utility for visualizing matrix data. Bioinformatics, 19, 295–6.

Payen, V., Chemin, A., Jonville-Bera, A. P., Saliba, E. & Cantagrel, S. 2006. [Fetal toxicity of angiotensin-II receptor antagonists]. J Gynecol Obstet Biol Reprod (Paris), 35, 729–31.

Perico, N., Askenazi, D., Cortinovis, M. & Remuzzi, G. 2018. Maternal and environmental risk factors for neonatal AKI and its long-term consequences. Nat Rev Nephrol, 14, 688-703. 10.1038/s41581-018-0054-y

Perner, B., Englert, C. & Bollig, F. 2007. The Wilms tumor genes wt1a and wt1b control different steps during formation of the zebrafish pronephros. Dev Biol, 309, 87-96. S0012-1606(07)01160-8 [pii] 10.1016/j.ydbio.2007.06.022

Piersma, A. H. 2004. Validation of alternative methods for developmental toxicity testing. Toxicol Lett, 149, 147-53. 10.1016/j.toxlet.2003.12.029

Pursley, T. J., Blomquist, I. K., Abraham, J., Andersen, H. F. & Bartley, J. A. 1996. Fluconazole-induced congenital anomalies in three infants. Clin Infect Dis, 22, 336-40. 10.1093/clinids/22.2.336

Ramoz, L. L. & Patel-Shori, N. M. 2014. Recent changes in pregnancy and lactation labeling: retirement of risk categories. Pharmacotherapy, 34, 389-95. 10.1002/phar.1385

Rodriguez, M. M., Gomez, A. H., Abitbol, C. L., Chandar, J. J., Duara, S. & Zilleruelo, G. E. 2004. Histomorphometric analysis of postnatal glomerulogenesis in extremely preterm infants. Pediatr Dev Pathol, 7, 17-25. 10.1007/s10024-003-3029-2

Russell, W. M. 1995. The development of the three Rs concept. Altern Lab Anim, 23, 298–304.

Sasagawa, S., Nishimura, Y., Kon, T., Yamanaka, Y., Murakami, S., Ashikawa, Y., Yuge, M., Okabe, S., Kawaguchi, K., Kawase, R. & Tanaka, T. 2016. DNA Damage Response Is Involved in the Developmental Toxicity of Mebendazole in Zebrafish Retina. Front Pharmacol, 7, 57. 10.3389/fphar.2016.00057

Schindelin, J., Arganda-Carreras, I., Frise, E., Kaynig, V., Longair, M., Pietzsch, T., Preibisch, S., Rueden, C., Saalfeld, S., Schmid, B., Tinevez, J. Y., White, D. J., Hartenstein, V., Eliceiri, K., Tomancak, P. & Cardona, A. 2012. Fiji: an open-source platform for biological-image analysis. Nat Methods, 9, 676-82. nmeth.2019 [pii] 10.1038/nmeth.2019

Singh, R. R., Cullen-Mcewen, L. A., Kett, M. M., Boon, W. M., Dowling, J., Bertram, J. F. & Moritz, K. M. 2007. Prenatal corticosterone exposure results in altered AT1/AT2, nephron deficit and hypertension in the rat offspring. J Physiol, 579, 503-13. 10.1113/jphysiol.2006.125773

Tsai, E. C., Brown, J. A., Veldee, M. Y., Anderson, G. J., Chait, A. & Brunzell, J. D. 2004. Potential of essential fatty acid deficiency with extremely low fat diet in lipoprotein lipase deficiency during pregnancy: A case report. BMC Pregnancy Childbirth, 4, 27. 10.1186/1471-2393-4-27

Werler, M. M., Mitchell, A. A., Hernandez-Diaz, S. & Honein, M. A. 2005. Use of over-the-counter medications during pregnancy. Am J Obstet Gynecol, 193, 771-7. 10.1016/j.ajog.2005.02.100

Westerfield, M. 2000a. The zebrafish book. A guide for the laboratory use of zebrafish (Danio rerio), 4th edition., University of Oregon Press, Eugene, OR.

Westerfield, M. 2000b. The zebrafish book. A guide for the laboratory use of zebrafish (Danio rerio). 4th ed., Univ. of Oregon Press, Eugene.

Westhoff, J. H., Giselbrecht, S., Schmidts, M., Schindler, S., Beales, P. L., Tonshoff, B., Liebel, U. & Gehrig, J. 2013. Development of an automated imaging pipeline for the analysis of the zebrafish larval kidney. PLoS One, 8, e82137. 10.1371/journal.pone.0082137 PONE-D-13-30412 [pii]

Wittbrodt, J. N., Liebel, U. & Gehrig, J. 2014. Generation of orientation tools for automated zebrafish screening assays using desktop 3D printing. BMC Biotechnol, 14, 36. 10.1186/1472-6750-14-36

Wong, B., Ooi, T. C. & Keely, E. 2015. Severe gestational hypertriglyceridemia: A practical approach for clinicians. Obstet Med, 8, 158-67. 10.1177/1753495x15594082

Xiao, W. L., Liu, X. Y., Liu, Y. S., Zhang, D. Z. & Xue, L. F. 2017. The relationship between maternal corticosteroid use and orofacial clefts-a meta-analysis. Reprod Toxicol, 69, 99-105. 10.1016/j.reprotox.2017.02.006

Zarn, J. A., Bruschweiler, B. J. & Schlatter, J. R. 2003. Azole fungicides affect mammalian steroidogenesis by inhibiting sterol 14 alpha-demethylase and aromatase. Environ Health Perspect, 111, 255-61. 10.1289/ehp.5785

