## Supplementary Tables for "In vivo high-content screening in zebrafish for developmental nephrotoxicity of approved drugs": Suppl Table 3.pdf

**Table S3. Main ATC D-level groups provoking renal developmental toxicity.**

| ATC D-level description | ATC code D-level | Chemical name |
|---|---|---|
| **Dihydropyridine derivatives** | C08CA | Amlodipine |
| | | **Felodipine** |
| | | **Isradipine** |
| | | Nicardipine hydrochloride |
| | | **Nifedipine** |
| | | Nimodipine |
| | | **Nitrendipine** |
| | | **Lacidipine** |
| | | **Nilvadipine** |
| | | **Cilnidipine** |
| | | Benidipine hydrochloride |
| **HMG CoA reductase inhibitors** | C10AA | **Simvastatin** |
| | | **Lovastatin** |
| | | Pravastatin |
| | | **Fluvastatin sodium salt** |
| | | **Atorvastatin** |
| **Fibrates** | C10AB | **Clofibrate** |
| | | Bezafibrate |
| | | **Gemfibrozil** |
| | | **Fenofibrate** |
| | | Ciprofibrate |
| **Imidazole and triazole derivatives** | D01AC | **Sulconazole nitrate** |
| | | Bifonazole |
| | | **Oxiconazole nitrate** |
| | | **Sertaconazole nitrate** |
| **Corticosteroids, moderately potent (group II)** | D07AB | **Flumethasone** |
| | | Flumethasone pivalate |
| | | **Desonide** |
| | | **Alclometasone dipropionate** |
| | | **Clocortolone pivalate** |
| **Corticosteroids, potent (group III)** | D07AC | **Fluocinonide** |
| | | **Fluocinolone acetonide** |
| | | Flurandrenolide |
| | | **Diflorasone diaceate** |
| | | Amcinonide |
| | | Prednicarbate |
| **Imidazole derivatives** | G01AF | Clotrimazole |
| | | Econazole nitrate |
| | | **Isoconazole** |
| | | **Tioconazole** |
| | | **Butoconazole nitrate** |

| | | |
|---|---|---|
| **Glucocorticoids** | H02AB | **Betamethasone** |
| | | **Dexamethasone acetate** |
| | | **Methylprednisolone, 6-alpha** |
| | | **Prednisolone** |
| | | Prednisone |
| | | **Triamcinolone** |
| | | **Hydrocortisone base** |
| | | **Cortisol acetate** |
| | | Cortisone |
| | | Rimexolone |
| | | Deflazacort |
| **Acetic acid derivatives and related substances** | M01AB | **Indometacin** |
| | | Sulindac |
| | | Tolmetin sodium salt dihydrate |
| | | **Zomepirac sodium salt** |
| | | **Diclofenac sodium** |
| | | Etodolac |
| | | **Fentiazac** |
| | | Acemetacin |
| | | **Ketorolac tromethamine** |
| | | **Aceclofenac** |
| | | Bufexamac |
| **Propionic acid derivatives** | M01AE | **S-(+)-ibuprofen** |
| | | **Naproxen** |
| | | Ketoprofen |
| | | **Fenoprofen calcium salt dihydrate** |
| | | **Fenbufen** |
| | | Suprofen |
| | | **Flurbiprofen** |
| | | **Indoprofen** |
| | | Tiaprofenic acid |
| | | **Oxaprozin** |
| **Benzimidazole derivatives** | P02CA | **Mebendazole** |
| | | Tiabendazole |
| | | **Albendazole** |
| | | Flubendazole |
| | | **Fenbendazole** |

Drugs affecting renal development as evaluated by quantitative measurements are highlighted in bold. Abbreviation: ATC, Anatomical Therapeutic Chemical Classification system.
