## Supplementary Tables for "In vivo high-content screening in zebrafish for developmental nephrotoxicity of approved drugs": Suppl Table 4.pdf

**Table 2. Additional compounds with adverse effects on pronephros development.**

| E-level description | ATC D-level description | ATC code D-level |
|---|---|---|
| Omeprazole | Proton pump inhibitors | A02BC |
| Dicyclomine hydrochloride | Synthetic anticholinergics, quaternary ammonium compounds | A03AA |
| Warfarin | Vitamin K antagonists | B01AA |
| **Proscillaridin A** | **Scilla glycosides** | **C01AB** |
| **Amiodarone hydrochloride** | **Antiarrhythmics, class III** | **C01BD** |
| Metaraminol bitartrate | Adrenergic and dopaminergic agents | C01CA |
| Midodrine hydrochloride | Adrenergic and dopaminergic agents | C01CA |
| Canrenone | Aldosterone antagonists | C03DA |
| Pentoxifylline | Purine derivatives | C04AD |
| **Suloctidil** | **Other peripheral vasodilators** | **C04AX** |
| **Diltiazem hydrochloride** | **Phenylalkylamine derivates** | **C08DB** |
| **Lidoflazine** | **Other non-selective calcium channel blockers** | **C08EX** |
| Valsartan | Angiotensin II antagonists, plain | C09CA |
| **Irbesartan** | **Angiotensin II antagonists, plain** | **C09CA** |
| **Ciclopirox ethanolamine** | **Other antifungals for topical use** | **D01AE** |
| **Isotretinoin** | **Retinoids for treatment of acne** | **D10BA** |
| **Norgestimate** | **Progestogens and estrogens, fixed combinations** | **G03AA** |
| **Progesterone** | **Pregnen (4) derivatives** | **G03DA** |
| **Danazol** | **Antigonadotropins and similar agents** | **G03XA** |
| Gestrinone | Antigonadotropins and similar agents | G03XA |
| **Fludrocortisone acetate** | **Mineralocorticoids** | **H02AA** |
| Methicillin sodium | Beta-lactamase resistant penicillins | J01CF |
| Sulbactam | Beta-lactamase inhibitors | J01CG |
| Tazobactam | Beta-lactamase inhibitors | J01CG |
| Cefotaxime sodium salt | Third-generation cephalosporins | J01DD |
| Troleandomycin | Macrolides | J01FA |
| Streptomycin sulfate | Streptomycins | J01GA |
| **Nalidixic acid sodium salt** | **Other quinolones** | **J01MB** |
| Oxolinic acid | Other quinolones | J01MB |
| **Amphotericin B** | **Antibiotics** | **J02AA** |
| **Miconazole** | **Imidazole derivates** | **J02AB** |
| Dapsone | Drugs for treatment of lepra | J04BA |
| Delavirdine | Non-nucleoside reverse transcriptase inhibitors | J05AG |
| Raltitrexed | Folic acid analogues | L01BA |
| **Carmofur** | **Pyrimidine analogues** | **L01BC** |
| Epirubicin hydrochloride | Anthracyclines and related substances | L01DB |
| Topotecan | Other antineoplastic agents | L01XX |
| **Flutamide** | **Anti-androgens** | **L02BB** |
| **Nilutamide** | **Anti-androgens** | **L02BB** |

| | | |
|---|---|---|
| Exemestane | Aromatase inhibitors | L02BG |
| **Leflunomide** | **Selective immunosuppressants** | **L04AA** |
| Azathioprine | Other immunosuppressants | L04AX |
| Thalidomide | Other immunosuppressants | L04AX |
| **Phenylbutazone** | **Butylpyrazolidines** | **M01AA** |
| **Piroxicam** | **Oxicams** | **M01AC** |
| **Meloxicam** | **Oxicams** | **M01AC** |
| **Mefenamic acid** | **Fenamates** | **M01AG** |
| Nimesulide | Other anti-inflammatory and antirheumatic agents, non-steroids | M01AX |
| Auranofin | Gold preparations | M01CB |
| **Etofenamate** | **Antiinflammatory preparations, non-steroids for topical use** | **M02AA** |
| **Felbinac** | **Antiinflammatory preparations, non-steroids for topical use** | **M02AA** |
| Cisatracurium besylate | Other quaternary ammonium compounds | M03AC |
| **Diflunisal** | **Salicylic acid and derivatives** | **N02BA** |
| Valproic acid | Fatty acid derivatives | N03AG |
| Entacapone | Other dopaminergic agents | N04BX |
| **Fluspirilen** | **Diphenylbutylpiperidine derivatives** | **N05AG** |
| **Pimozide** | **Diphenylbutylpiperidine derivatives** | **N05AG** |
| Asenapine maleate | Diazepines, oxazepines and thiazepines | N05AH |
| **Isocarboxazid** | **Monoamine oxidase inhibitors, non-selective** | **N06AF** |
| **Disulfiram** | **Drugs used in alcohol dependence** | **N07BB** |
| Isoflupredone acetate | n/a | n/a |
| **Nocodazole** | **n/a** | **n/a** |
| **Retinoic acid** | **n/a** | **n/a** |
| **GBR 12909 dihydrochloride** | **n/a** | **n/a** |
| Pregnenolone | n/a | n/a |
| Parthenolide | n/a | n/a |
| **Mevastatin** | **n/a** | **n/a** |
| Tranilast | n/a | n/a |
| Hexestrol | n/a | n/a |
| **Cycloheximide** | **n/a** | **n/a** |
| Butylparaben | n/a | n/a |
| **Methiazole** | **n/a** | **n/a** |
| Homosalate | n/a | n/a |
| **Clonixin Lysinate** | **n/a** | **n/a** |
| **Ethoxzolamide** | **n/a** | **n/a** |
| **Benzoxiquine** | **n/a** | **n/a** |
| **Clioquinol** | **Hydroxyquinoline derivatives** | **P01AA** |
| Diloxanide furoate | Dichloracetamide derivatives | P01AC |
| **Halofantrine hydrochloride** | **Other antimalarials** | **P01BX** |
| Ivermectin | Avermectines | P02CF |
| Altrenogest | n/a | n/a |

| | | |
|---|---|---|
| **Flunixin meglumine** | **n/a** | **n/a** |
| **Oxibendazole** | **n/a** | **n/a** |
| **Parbendazole** | **n/a** | **n/a** |
| Flunisolide | Corticosteroids | R01AD |
| Mometasone furoate | Corticosteroids | R01AD |
| **Salmeterol** | **Selective beta-2-adrenoreceptor agonists** | **R03AC** |
| Beclomethasone dipropionate | Glucocorticoids | R03BA |
| Budesonide | Glucocorticoids | R03BA |
| Fluticasone propionate | Glucocorticoids | R03BA |
| Meclozine dihydrochloride | Piperazine derivatives | R06AE |
| Astemizole | Other anthistamines for systemic use | R06AX |

Drugs with a strong effect on pronephros development as judged by quantitative analysis are highlighted in bold. Abbreviation: ATC, Anatomical Therapeutic Chemical Classification system.
