## Supplementary Figures for "In vivo high-content screening in zebrafish for developmental nephrotoxicity of approved drugs"

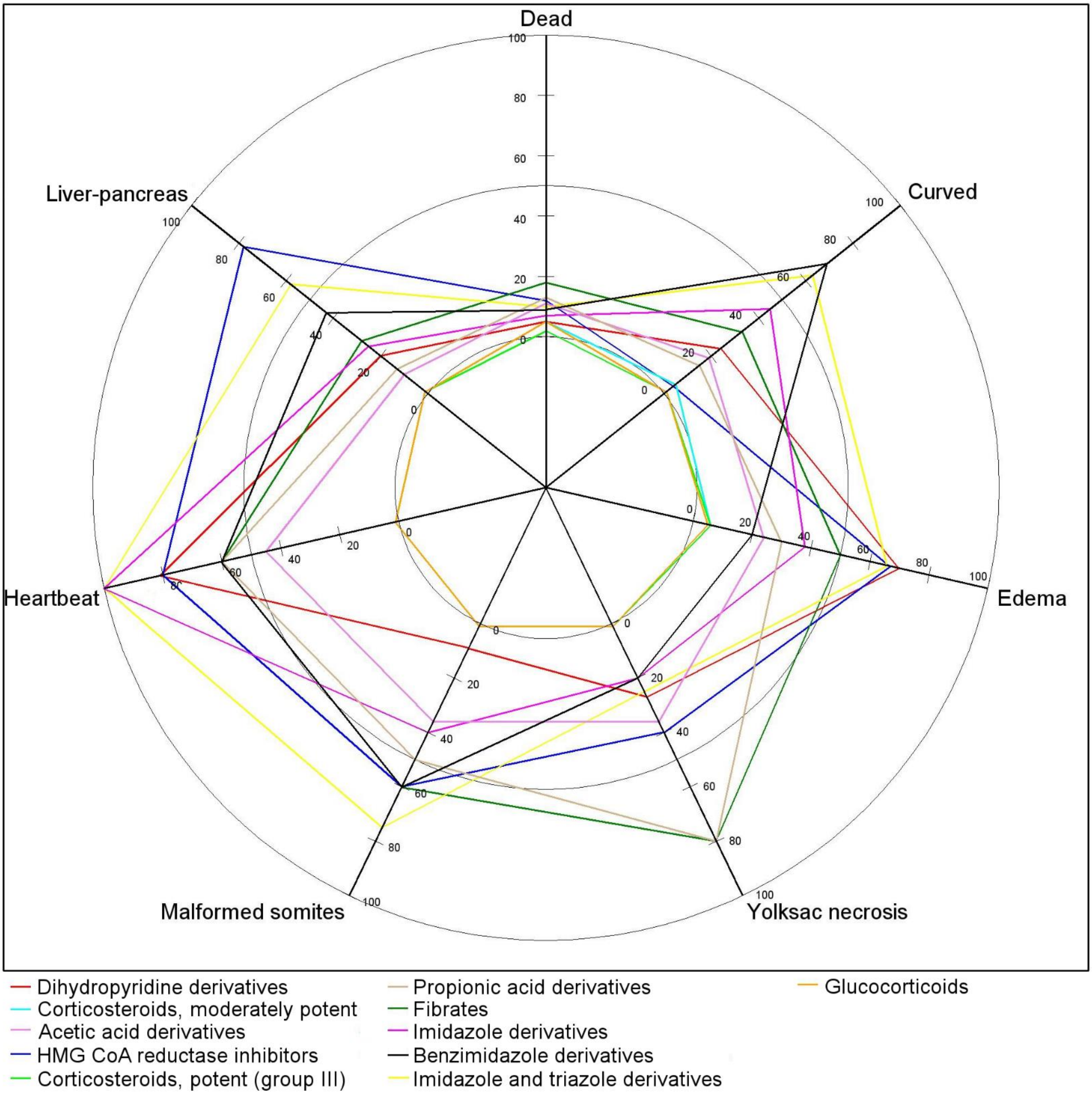


**Supplementary Figure 1| Extrarenal phenotypes of larval zebrafish following exposure to eleven nephrotoxic ATC code D-Level substance groups during development.** Radar plot illustrating percentage of positive larval zebrafish for the following parameters: lethality (dead), curved back (curved), pericardial edema (edema), yolk sac necrosis, malformed somites, heartbeat alteration (heartbeat), loss of liver-pancreas fluorescence (liver-pancreas). Analysis by evaluation of 20 larvae per substance. Line color description as indicated.

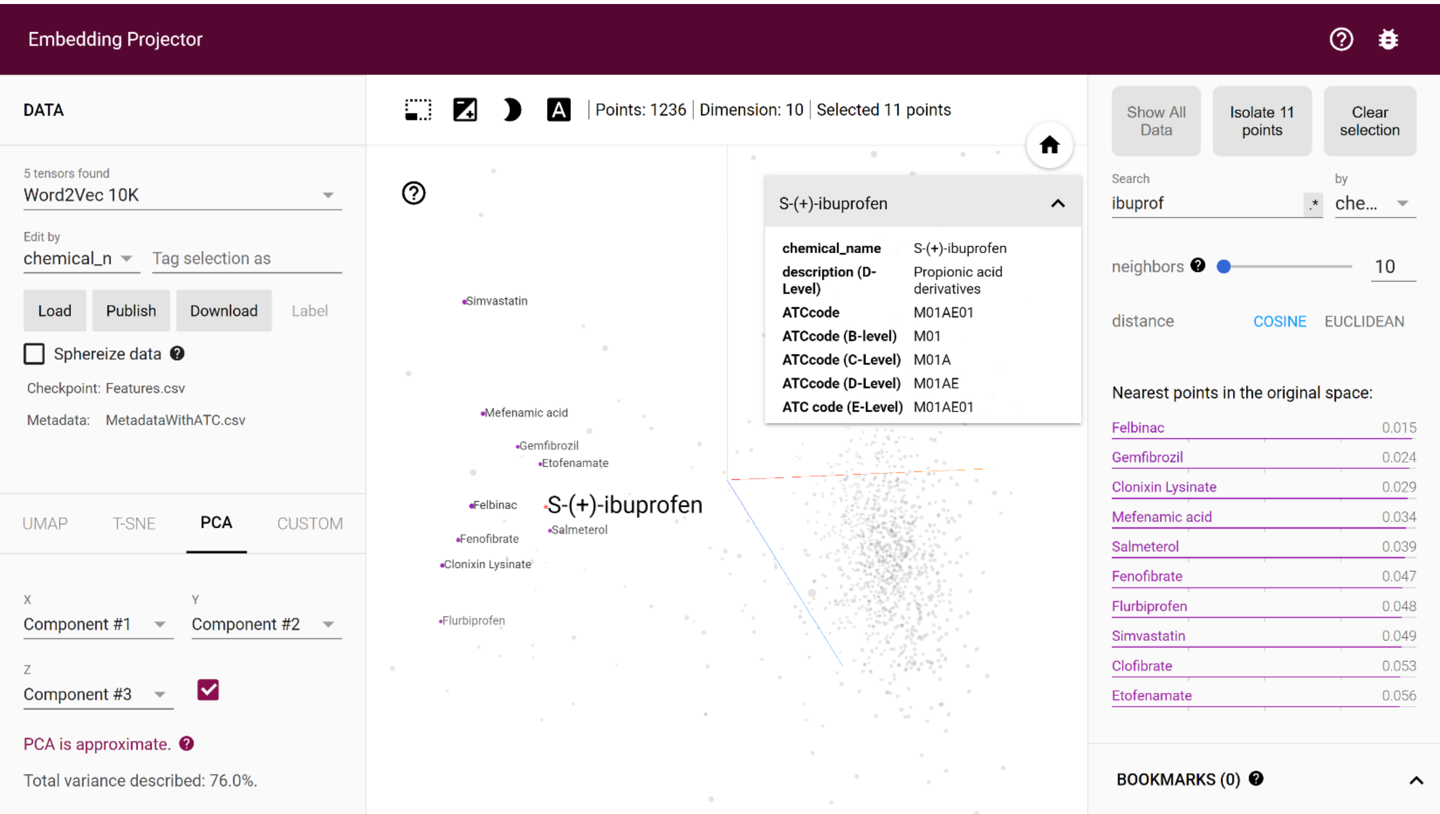


**Supplementary Figure 2| Screenshot of a view of the dataset in the tensorflow-projector interactive web-interface.** The interface (https://projector.tensorflow.org/) performs dimensionality reduction down to 3 dimensions using principal component analysis (PCA). Shown are PCA values calculated on z-scores for the morphological features as in Figure 3 and Supplementary Table 1. Additional modes of projection are available (UMAP and T-SNE, not shown). Each point in the cloud corresponds to a tested compound. Compounds with similar morphological features end up in the same neighbourhood of the PCA projection. In this example, the compound S-(+)-ibuprofen is selected in red and the 10 nearest-neighbour compounds are highlighted in violet, and listed with their distance to the selected compound on the right panel. The distance between compounds is computed as the cosine between the vectors made of the morphological features for each compound (feature vectors).

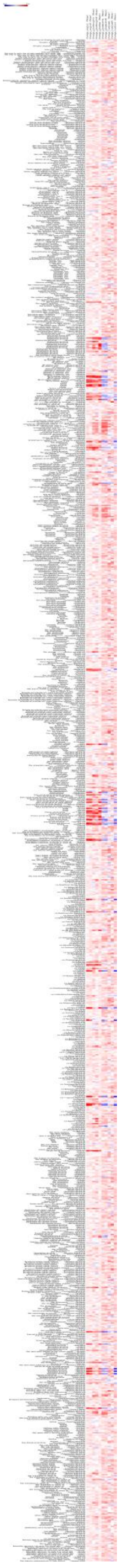

Note: This is a thumbnail only. Please download a fully annotated heatmap (737 x 10127 pixels ) from Zenodo (click link).

**Supplementary Figure 3| Fully annotated heatmap of quantitative features.** Shown are z-scores of quantitative morphological measurements. For legend please refer to Figure 3 of the main manuscript.

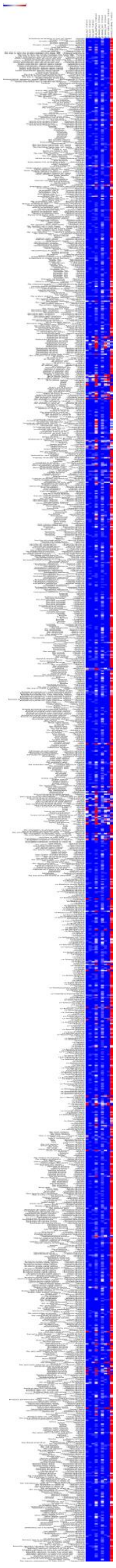

Note: This is a thumbnail only. Please download a fully annotated heatmap (737 x 10119 pixels ) from Zenodo (click link).

**Supplementary Figure 4| Fully annotated heatmap of qualitative features.** Shown are ratios of embryos assigned with a quantitative category using the MAT tool. For legend please refer to Figure 3 of the main manuscript.
